# NatB-dependent acetylation protects procaspase-8 from UBR4-mediated degradation and is required for full induction of the extrinsic apoptosis pathway

**DOI:** 10.1101/2023.09.25.559278

**Authors:** Joana P. Guedes, Jean Baptiste Boyer, Jasmine Elurbide, Beatriz Carte, Virginie Redeker, Laila Sago, Thierry Meinnel, Manuela Côrte-Real, Carmela Giglione, Rafael Aldabe

## Abstract

N-terminal acetyltransferase B (NatB) is a major contributor to the N-terminal acetylome and is implicated in several key cellular processes including apoptosis and proteostasis. However, the molecular mechanisms linking NatB-mediated N-terminal acetylation to apoptosis and its relationship with protein homeostasis remain elusive. In this study, we generated mouse embryonic fibroblasts (MEFs) with an inactivated catalytic subunit of NatB (*Naa20*^-/-^) to investigate the impact of NatB deficiency on apoptosis regulation. Through quantitative N-terminomics, label-free quantification, and targeted proteomics, we demonstrated that NatB does not influence the proteostasis of all its substrates. Instead, our focus on putative NatB-dependent apoptotic factors revealed that NatB-mediated acetylation serves as a protective shield against UBR4 and UBR1 Arg/N-recognin-mediated degradation. Notably, *Naa20*^-/-^ MEFs exhibited reduced responsiveness to extrinsic pro-apoptotic stimuli, a phenotype that was partially reversible upon UBR4 Arg/N-recognin silencing and consequent inhibition of procaspase-8 degradation. Collectively, our results shed light on how the interplay between NatB-mediated acetylation and the Arg/N-degron pathway impacts apoptosis regulation, providing new perspectives in the field including in therapeutic interventions.

## INTRODUCTION

N-terminal protein modifications (NPMs) are among the first and fastest responses to the cellular environment, influencing not only the modified protein but also cell, tissue, and whole organism phenotypes^1, 2, 3, 4^. One major NPM is N-terminal (Nt-) acetylation, an essential multitasking protein modification implicated in many diseases^5, 6^, where an acetyl group is transferred from acetyl coenzyme A to the α-amino group of a protein N-terminus catalyzed by Nt-acetyltransferases (NATs)^5, 7^. The eight currently known eukaryotic NATs have been defined based on their subunits and substrate specificity^6, 8^. Nt-acetyltransferase B (NatB) comprises the catalytic subunit NAA20 and the auxiliary subunit NAA25. NAA25 binds to the ribosome and ensures co-translational activity of NatB^9^, while the catalytic site of NAA20 anchors and acetylates the N-alpha amino group of the initiator methionine (iMet) of all proteins immediately followed by an asparagine, glutamine, aspartic or glutamic acid residue^10, 11^.Together with NatA, which acetylates proteins that have lost their iMet, NatB is the major contributor to Nt-acetylation. NatB substrates cover nearly 21% of human, *Caenorhabditis elegans*, and *Arabidopsis thaliana* proteomes and 15% of yeast proteome. In all these organisms, almost all NatB substrates are fully acetylated, unlike the substrates of other NATs (especially NatA), many of which are only partially modified^10, 12, 13^.

For a few well-studied proteins, NatB has been shown to be involved in promoting resistance to aggregation, protein-protein interactions, protein sorting to distinct cellular compartments, and regulating protein half-life^6, 14^. However, the overall impact of NatB-mediated Nt-acetylation on proteostasis is still poorly understood. Global protein stability does not seem to be affected by NatB subunit depletion in yeast or human cells^15, 16, 17^, suggesting that NatB-mediated Nt-acetylation may not be a dominant factor in proteostasis. Conversely, several studies have highlighted the importance of NATs-mediated Nt-acetylation on the homeostasis of specific proteins in the context of N-degrons, which are N-terminal degradation signals recognized by the N-degron pathways^18^. For instance, the N-terminal acetylated variants of MX-Rgs2, a G-protein regulator, have been shown to be selectively targeted by the Ac/N-end rule pathway when X is either Arg or Glu^19^.

Whatever the impact of NatB-mediated Nt-acetylation on proteostasis or protein half-life, the inactivation or downregulation of any NatB subunit influences fundamental cellular processes such as actin cytoskeleton organization^10, 20–23^, NAD^+^ homeostasis^24^, influenza virus protein PA-X ^25^, proliferation^22, 23, 26^, and apoptosis^23, 26, 27^. However, the exact molecular mechanisms linking defective NatB or NatB-mediated Nt-acetylation to specific cellular processes and their interdependence have not yet been elucidated.

Given the prevalence and importance of Nt-acetylation and apoptosis in numerous human diseases^6, 28^, we sought to establish the mechanistic role of NatB-dependent Nt-acetylation on apoptosis regulation. Here, we show that NatB inactivation in mouse embryonic fibroblasts (MEFs) leads to degradation of proapoptotic proteins, particularly of procaspase-8, mediated by the UBR4 E3 ubiquitin ligase, thereby limiting activation of the extrinsic apoptotic pathway. Our findings provide the first evidence implicating NatB-Arg/N-degron pathway in apoptosis regulation.

## METHODS

### Standard methods are reported in the Supplementary Information Inactivation of *Naa20* in MEFs

MEFs were obtained from *Naa20*^tm1a(KOMP)Wtsi^ transgenic mice as described in the **Supplementary Methods**. *Naa20* was inactivated with a recombinant adenovirus expressing CRE recombinase (Ad5CMVCre, MOI 2000). An empty adenovirus was used as a negative control (AdEmpty, MOI 2000). Recombinant adenovirus was inoculated in DMEM supplemented with 2% FBS and 1% penicillin/streptomycin and, 24 h later, regular DMEM was added. Cells were trypsinized and re-plated, two or five days post-infection.

### Apoptosis assays

Cells were seeded in six-well plates, infected, and six days post-infection treated with 50 µM etoposide, 100 ng/mL TNF-α plus 500 nM SMAC, 5 µM of MG132, or 5 ng/µL of tunicamycin for 0, 8, 12, and 24 h. At each time point, cells were harvested and collected for protein extraction for further analysis by western blotting.

### N-terminal acetylation assays

Eight biological replicates of WT or *Naa20*^-/-^ MEFs (10^6^ cells) were harvested and six days after adenovirus infection with AdEmpty and Ad5CMVCre, respectively, were centrifuged at 6000 rpm for 5 min at 4°C, and pellets were washed with PBS 1× and centrifuged again. Next, cell pellets were resuspended in 300 µL of protein extraction buffer (50 mM HEPES/NaOH pH 7.2; 1.5 mM MgCl_2_; 1 mM EGTA; 10% glycerol; 1% Triton X-100; 150 mM NaCl; 2 mM PMSF; 1 protease inhibitor cocktail tablet (EDTA+) in 50 mL), subjected to lysis by sonication, centrifuged at 17 000 rpm for 20 min at 4°C, and the supernatant recovered for proteomic analysis.

For each sample, 1 mg of protein was used following the SILProNAQ protocol previously described^29^. Briefly, N-acetoxy-[^2^H_3_] succinimide was used to label protein N-termini and lysine ε-amino groups before being digested by trypsin. Peptides were then fractionated by chromatography using a strong cation exchange (SCX) column to discriminate acetylated and unacetylated internal peptides. Individual fractions 2-11 were analyzed by LC-MS using 55 min methods on the LTQ-Orbitrap Velos mass spectrometer (Thermo Fisher Scientific, Waltham, MA), and three pools of fractions (2-5, 6-8, and 9-11) were analyzed using 100 min methods on the TIMS-ToF (Bruker, Billerica, MA). Data were searched against the *Mus musculus* SwissProt database through Mascot Distiller then parsed using the in-house eNcounter script to validate the detected N-termini and obtain the Nt-acetylation yield^30^.

### Label-free proteome quantitation

In parallel with the N-terminal acetylation assay, the same MEF samples were used for full proteomic analysis. 30 µg of protein extract were loaded on an SDS-PAGE gel and digested following an in-gel digestion protocol^31^. The resulting peptides were extracted, dried, and then analyzed on the TIMS-ToF mass spectrometer using the same 100 min method as before. Data were processed using MaxQuant to identify and quantify.

### siRNA silencing of E3 ubiquitin ligases

WT or *Naa20*^-/-^ MEF cells were seeded in six-well plates and infected the day after with the corresponding adenovirus as described above. Two days later, cells were trypsinized and re-plated as described previously and, four days post-infection, cells were re-plated and transfected with the corresponding siRNAs. 150 000 cells/ml were transfected in suspension with the siRNAs (**Supplementary Methods**) and the Lipofectamine^®^ RNAiMAX Transfection Reagent (Invitrogen, Thermo Fisher Scientific 13778) according to manufacturer’s instructions. Transfected cells were collected 48 h post-transfection for western blot and real-time PCR analysis.

### Statistical analysis

Statistical analyses were performed using Prism 8 (GraphPad Software, La Jolla, CA). Normality of groups was assessed with the Shapiro-Wilk test. The unpaired *t*-test was used to compare parametric data the Mann-Whitney test for non-parametric data. Two-way ANOVA was used to analyze proliferation data. The significance of enriched NatB substrates in different MS quantifications was calculated using the built-in two-tailed *t*-test function for comparison of two samples with equivalent variances in Microsoft Excel for the N-terminome analysis, while a two sample Student *t*-test, with a threshold p-value of 0.05, was used in Perseus for the label-free proteome analysis.

## RESULTS

### Complete *Naa20* inactivation induces cytoskeletal abnormalities and decreases proliferation

Based on our previous studies that showed increased sensitivity of *Naa25^-/-^* MEFs ^27^ and *Naa20^-/-^* HeLa cells^26^ to the apoptotic inducer MG132, we aimed to further investigate the role of NatB acetylation on apoptosis by inactivating the NatB catalytic subunit in MEFs.

Six days of infection with Ad5CMVCre (*Naa20*^-/-^ MEFs) resulted in a 99.6% reduction of *Naa20* transcript compared with AdEmpty-infected MEFs (WT) (**Fig. 1A**), and NAA20 protein was no longer detectable in *Naa20*^-/-^ MEFs (**Fig. 1B**). *Naa20* inactivation was associated with a significant reduction in *Naa20*^-/-^ MEF proliferation four days after infection (**Fig. 1C**). Moreover, six days after AdCre infection, actin fibers became disorganized in *Naa20*^-/-^ MEFs compared to WT (**Fig. 1D**), and, in keeping with this actin disorganization, focal adhesions decreased, a phenotype commonly observed upon NatB complex inactivation or downregulated in other cell types. These results confirmed that *Naa20* is inactivated in *Naa20*^-/-^ MEF cells and induces phenotypes similar to those previously observed in other NatB-downregulated cellular contexts^10, 20, 21, 23^.

**Fig. 1.**
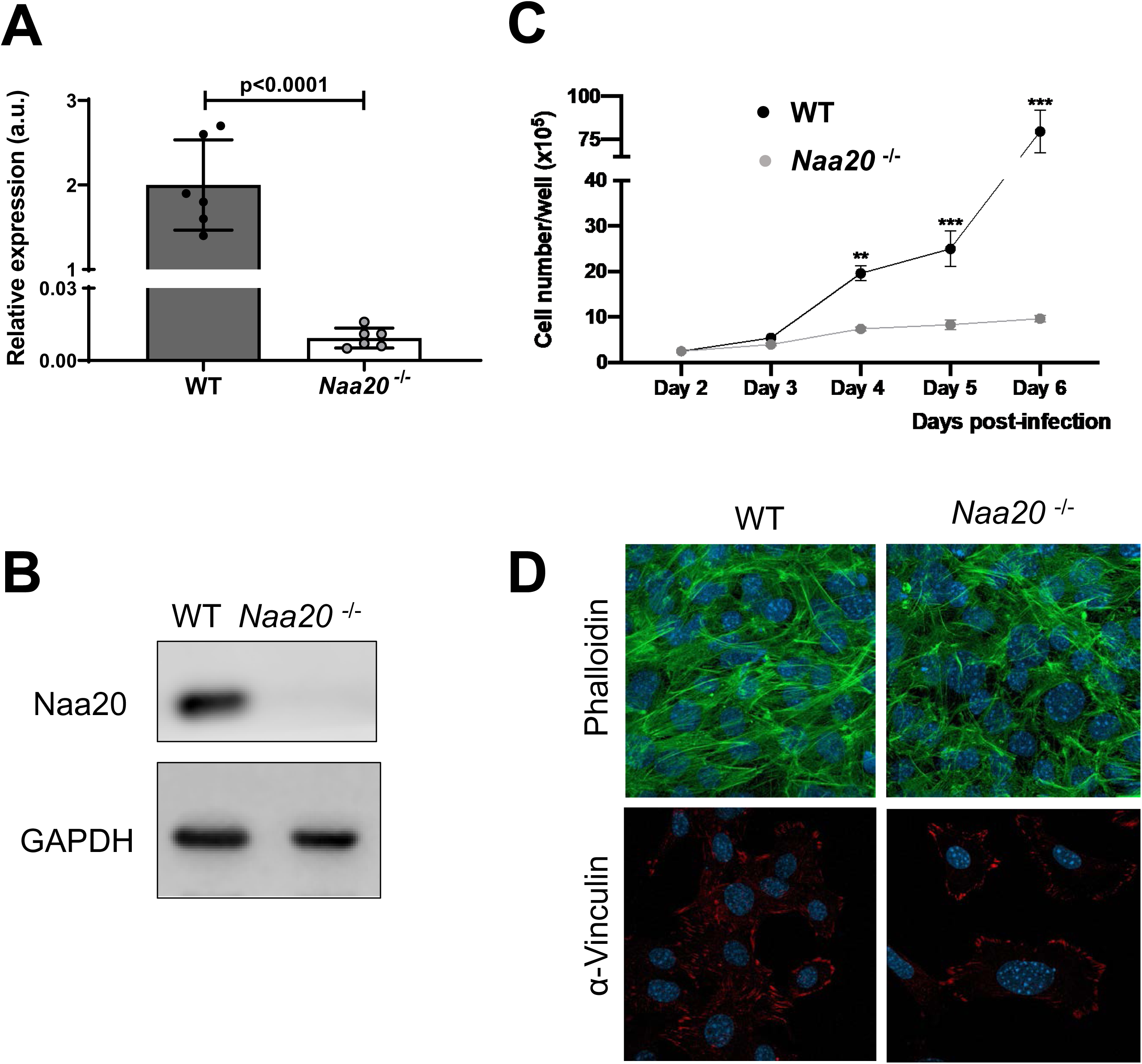
In *Naa20*^-/-^ MEFs lacking the catalytic subunit of NatB, *Naa20* mRNA and NAA20 protein are not expressed, and cellular proliferation and the actin cytoskeleton are perturbed. **(A)** *Naa20* mRNA quantification analysis in MEFs infected with the AdEmpty (WT) and Ad5CMVCre (*Naa20*^-/-^) by RT-PCR. Data are the mean ± SEM of six independent experiments and are normalized to a housekeeping gene (Histone 3) and analyzed by Student’s *t*-test. **(B)** NAA20 protein levels were detected by western blotting in WT and *Naa20*^-/-^cells using antibodies targeting NAA20. **(C)** Cell proliferation was visualized as the number of cells per well along post-infection days. Values represent the average number of WT or *Naa20*^-/-^ MEFs from three independent replicates as mean ± SEM (Student’s *t*-test). **(D)** Representative confocal microscopy images of actin (phalloidin) and focal adhesions (vinculin) in WT or *Naa20*^-/-^ MEFs.

### Quantitative N-terminomics of *Naa20*^-/-^ MEFs reveals reduction in Nt-acetylation of only NatB substrates

We next performed N-terminomics analysis of WT and *Naa20*^-/-^ MEFs using the ‘<u>S</u>table Isotope <u>L</u>abeling <u>Pro</u>tein <u>N</u>-terminal <u>A</u>cetylation <u>Q</u>uantification’ method to identify and quantify N-termini (SILProNAQ^29^). Analysis of WT and *Naa20*^-/-^ MEF cells identified 25,048 protein N-termini corresponding to 1988 non-redundant proteoforms (**Supplementary Dataset 1 and Supplementary Table 1**). The acetylation yields of 1191 of them (1073 in the WT and 945 in the NatB mutant background) could be quantified. The analysis of these unique proteoforms unravelled that 769 underwent removal of the initial Methionine (iMet) in agreement with known N-terminal methionine excision (NME) rules in eukaryotes ^32^, whereas 258 proteoforms retained their iMet **(Supplementary Table 1)**. The remaining 164 underwent larger cleavages in keeping with leader peptide removal. The quasi-totality of the iMet starting N-termini featured at position two an amino acid with a large lateral chain (**Supplementary Dataset 1**).

In *Naa20*^-/-^ MEFs, there was an overall reduction in Nt-acetylation, primarily in proteins retaining their iMet (**Fig. 2A, B**), with a relative decrease in the number of fully acetylated N-termini to only partially acetylated N-termini (**Supplementary Dataset 1** and **Supplementary Table 1**). By contrast, acetylation of N-termini devoid of iMet or N-termini processed downstream were unaffected (**Fig. 2C**, **D**). We compared our N-terminomics data to those previously reported for human cells in which NatB was downregulated^10^. Common identified and quantified N-termini of the two datasets revealed mostly decreases of the acetylation yields of NatB substrates (**Supplementary Dataset 2**). This effect was significantly stronger in *Naa20*^-/-^ MEFs (**Supplementary Dataset 2**). N-termini with the highest decreased Nt-acetylation in *Naa20*^-/-^ MEFs (26, **Table 1** and **Fig. 2E, F**) corresponded to NatB substrates (23), of which 18 displayed Asp and Glu at position two and, to a minor extent (5), Asn and Gln (**Table 1** and **Supplementary Dataset 1**). In *Naa20*^-/-^ MEFs, only three non-NatB substrates also showed strongly decreased Nt-acetylation (**Table 1**), but for which no information is available for the human counterparts (**Supplementary Dataset 2**).

**Fig. 2.**
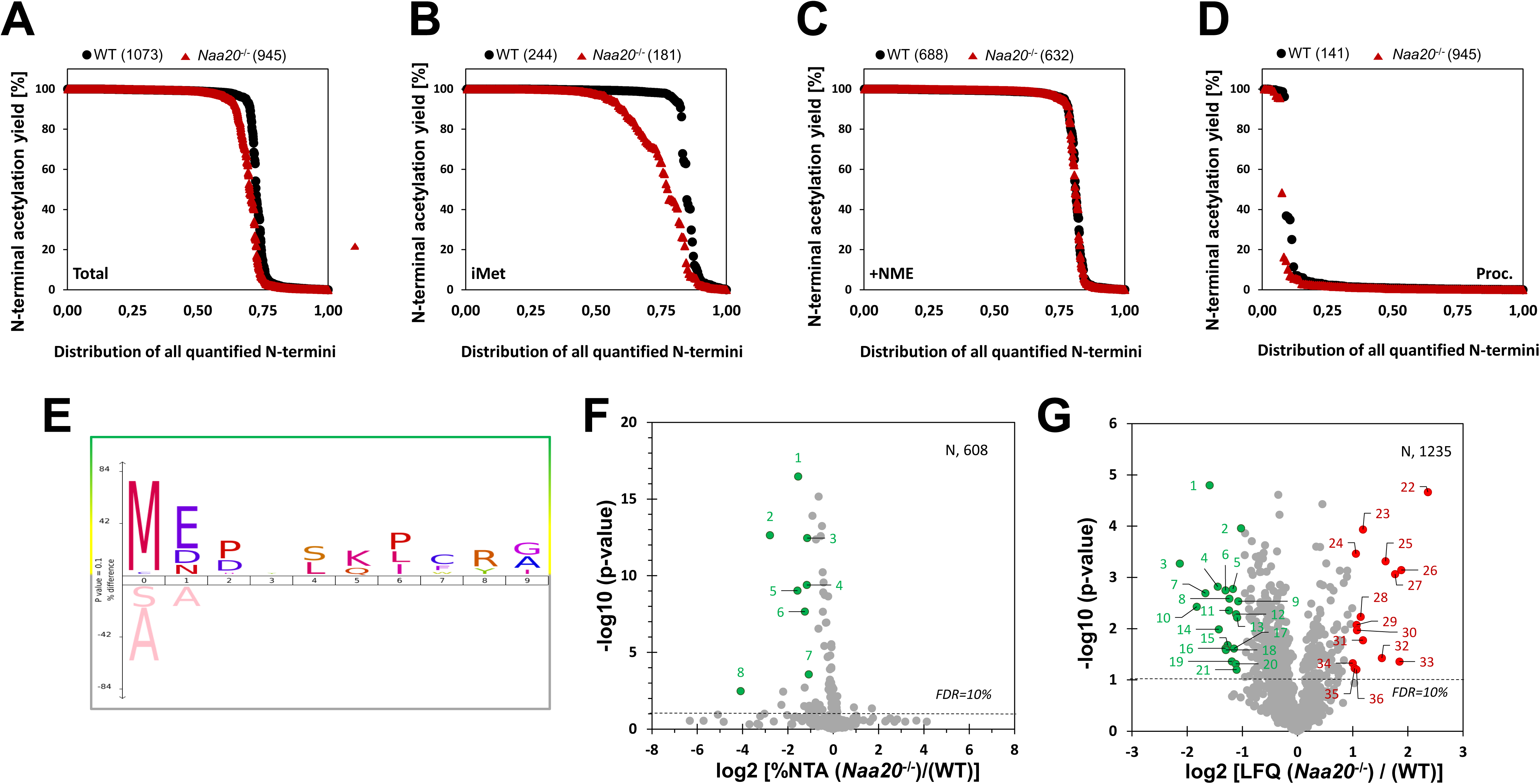
N-terminomic and proteomic analyses of *Naa20*^-/-^ MEFs reveal reduced N-terminal (Nt-) acetylation, specifically of NatB substrates, and deregulated protein levels. Accumulated distribution representing the percentage of Nt-acetylated **(A)** total proteome, **(B)** peptides with the initial methionine (iMet) present, **(C)** peptides with Nt-methionine excised, and **(D)** peptides with processed N-termini identified in normal (WT) and *Naa20*^-/-^ MEFs. **(E)** IceLogo representation of the N-terminal sequences with a minimum 20% decrease in NTA yield to those with <5% variation, when comparing the *Naa20*^-/-^ to the WT MEFs. Volcano plot representing **(F)** proteins with an Nt-acetylation reduction of 40% or more (green spots). The dashed horizontal line shows the p-value cut off, and the eight points highlighted in green indicate the affected proteins with the highest statistical significance. **(G)** Proteins with a label-free quantification (LFQ protein ratio) either <0.5 (green spots) or >2.0 (red spots) in the *Naa20*^-/-^ compared with WT samples. The dashed horizontal line shows the p-value cut off, and the points highlighted in green and red represent the downregulated and upregulated proteins, respectively. All plots contain only the N-termini that have been quantified at least once in each condition (WT or *Naa20*^-/-^). FDR - false discovery rate; N – number of samples.

**Table 1.** Most affected N-terminal acetylated substrates resulting from *Naa20* knockout in MEFs and their correlation with differences in protein expression when label-free quantitation was possible in WT and mutant samples. Proteins with a significant decrease in Nt-acetylation (NTA) according to a FDR <5%. NTA Pos.: NTA position; Nt-Seq.: N-terminal sequence; n.i.: protein not identified in the experiment; #: the first eight most statistically significant NTA downregulated proteins shown in Fig. 2F are highlighted in grey.

| Plot# | Entry Name | Protein Description | NTA Pos. | Nt- Seq. | %NTA (WT) | %NTA (Naa20<sup>-/-</sup>) | NTA Difference (Naa20<sup>-/-</sup> - (WT)) | NTA Ratio (Naa20<sup>-/-</sup>)/(WT) | LFQ Protein Ratio (Naa20<sup>-/-</sup>/WT) |
| --- | --- | --- | --- | --- | --- | --- | --- | --- | --- |
| 1 | RS24_MOUSE | 40S ribosomal protein S24 | 1 | MNDTVTIRTR | 99.4 ± 0.6 | 33.9 ± 3.4 | -65.5 | 0.341 | 0.693 |
| 2 | TM258_MOUSE | Oligosaccharyl transferase subunit TMEM258 | 1 | MELEAMSRYT | 93.3 ± 4.2 | 13.4 ± 2.1 | -79.9 | 0.144 | - |
| 3 | PRUN1_MOUSE | Exopolyphosphatase PRUNE1 | 1 | MEDYLQDCRA | 99.8 ± 0.3 | 44.9 ± 1.1 | -54.9 | 0.450 | 2.075 |
| 4 | IPO7_MOUSE | Importin-7 | 1 | MDPNTIIEAL | 99.5 ± 0.8 | 44.3 ± 3.5 | -55.1 | 0.446 | 0.516 |
| 5 | IFM3_MOUSE | Interferon-induced transmembrane protein 3 | 1 | MNHTSQAFIT | 97.9 ± 0.6 | 32.8 ± 1.8 | -65.2 | 0.335 | - |
| 6 | KEAP1_MOUSE | Kelch-like ECH-associated protein 1 | 1 | MQPEPKLSGA | 99.1 ± 0.7 | 41.5 ± 1.4 | -57.6 | 0.419 | - |
| 7 | TEBP_MOUSE | Prostaglandin E synthase 3 | 1 | MQPASAKWYD | 86.2 ± 7.3 | 40.7 ± 0.3 | -45.5 | 0.472 | 0.624 |
| 8 | AAAS_MOUSE | Aladin | 2 | CSLGLFPPPP | 99.9 ± 0.0 | 5.9 ± 5.5 | -94.1 | 0.059 | - |
| 9 | INT3_MOUSE | Integrator complex subunit 3 | 1 | MELQKGKGTV | 98.6 ± 0.9 | 26.1 | -72.5 | 0.265 | - |
| 10 | WDR46_MOUSE | WD repeat-containing protein 46 | 1 | METAPKPGRG | 100.0 | 50.3 ± 0.7 | -49.7 | 0.503 | - |
| 11 | PRAF3_MOUSE | PRA1 family protein 3 | 1 | MDVNLAPLRA | 99.7 ± 0.3 | 52.8 ± 3.8 | -46.9 | 0.529 | - |
| 12 | BABA2_MOUSE | BRISC and BRCA1-A complex member 2 | 1 | MSPEIALNRI | 99.2 ± 0.2 | 57.8 | -41.4 | 0.583 | - |
| 13 | RFA3_MOUSE | Replication protein A 14 kDa subunit | 1 | MEDIMQLPKA | 99.6 ± 0.4 | 58.5 ± 3.1 | -41.1 | 0.588 | - |
| 14 | ILK_MOUSE | Integrin-linked protein kinase | 1 | MDDIFTQCRE | 99.9 ± 0.1 | 63.4 ± 0.9 | -36.5 | 0.635 | - |
| 15 | GTR1_MOUSE | Solute carrier family 2 | 1 | MDPSSKKVTG | 99.3 ± 0.7 | 63.3 ± 2.3 | -36.0 | 0.638 | 0.623 |
| 16 | P4K2A_MOUSE | Phosphatidylinositol 4-kinase type 2-alpha | 1 | MDETSLVSP | 99.0 ± 0.8 | 66.7 | -32.3 | 0.674 | - |
| 17 | FBX22_MOUSE | F-box only protein 22 | 1 | MEPAGGGGV | 99.9 ± 0.0 | 67.9 ± 2.9 | -32.0 | 0.680 | - |
| 18 | TTL12_MOUSE | Tubulin--tyrosine ligase-like protein 12 | 1 | MEIQSGPQPG | 99.9 ± 0.1 | 70.7 | -29.2 | 0.708 | 0.748 |
| 19 | KT3K_MOUSE | Ketosamine-3-kinase | 1 | METLLKRELG | 99.9 ± 0.0 | 70.9 ± 0.1 | -29.1 | 0.709 | - |
| 20 | BCD1_MOUSE | Box C/D snoRNA protein 1 | 1 | MESAAEKEGT | 99.5 ± 0.5 | 71.9 ± 1.8 | -27.5 | 0.723 | - |
| 21 | NDUAC_MOUSE | NADH dehydrogenase 1 alpha subcomplex subunit 12 | 1 | MELVEVLKRG | 99.9 ± 0.0 | 73.3 ± 2.0 | -26.7 | 0.733 | - |
| 22 | FARP1_MOUSE | FERM, RhoGEF and pleckstrin domain-containing protein 1 | 2 | GEIEQKPTPA | 96.9 ± 2.7 | 70.4 ± 2.6 | -26.6 | 0.726 | 0.936 |
| 23 | TCPA_MOUSE | T-complex protein 1 subunit alpha | 1 | MEGPLSVFGD | 99.6 ± 0.5 | 74.0 ± 4.3 | -25.5 | 0.743 | 0.944 |
| 24 | AN32E_MOUSE | Acidic leucine-rich nuclear phosphoprotein 32 family member E | 1 | MEMKKKINME | 99.9 ± 0.0 | 75.8 ± 2.6 | -24.2 | 0.758 | 0.990 |
| 25 | THIC_MOUSE | Acetyl-CoA acetyltransferase | 1 | MNAGSDPVVI | 99.6 ± 0.3 | 76.9 ± 4.4 | -22.8 | 0.772 | 0.921 |
| 26 | SYRC_MOUSE | Arginine--tRNA ligase | 1 | MDGLVAQCSA | 99.5 ± 0.5 | 79.5 ± 4.7 | -20.0 | 0.799 | 1.009 |
|  | CASP3_MOUSE | Procaspase-3 | n.i. | n.i. | n.i. | n.i. | n.i. | n.i. | 0.649 |
|  | CASP8_MOUSE | Procaspase-8 | n.i. | n.i. | n.i. | n.i. | n.i. | n.i. | n.i. |
|  | CASP9_MOUSE | Procaspase-9 | n.i. | n.i. | n.i. | n.i. | n.i. | n.i. | n.i. |
|  | MARH6_MOUSE | E3 ubiquitin-protein ligase MARCHF6 | 1 | MDTAEEDICR | NTA | NTA | - | - | n.i. |
|  | UBR4_MOUSE | E3 ubiquitin-protein ligase UBR4 | n.i. | n.i. | n.i. | n.i. | n.i. | n.i. | 0.699 |

We observed that the functions of many retrieved proteins with reduced Nt-acetylation in *Naa20*^-/-^ MEFs were apoptosis-related (**Supplementary Dataset 1** and **Table 1**). Among them, we found TMEM258, a central mediator of ER quality control via apoptosis^33^ ; importin 7, which is involved via the Hippo pathway in the expression of genes important in apoptosis^34^; and IFM3, a member of the Interferon-induced transmembrane proteins (IFITMs) family^35^. This intriguing observation led us to note that procaspases-3, -8, -9, BAX, APAF1, and BID, which are major contributors to extrinsic and intrinsic apoptosis (**Supplementary** Fig.1), systematically displayed Asp or Glu at position 2. These proteins are also inferred as NatB substrates based on sequence similarity between mouse and human (100% conservation of the first amino acids) and a combination of experimental and computational evidence^10, 36–38^. Unfortunately, our SilProNAQ approach did not allow us to retrieve and analyze these N-termini due to the distal position of Arg residues required for trypsin cleavage and peptide identification, combined with low protein expression (**Table 1**).

To further explore the relationship between NatB-dependent Nt-acetylation and proteostasis, we next examined the NatB-dependent proteome, particularly that of caspases. Label-free quantification (LFQ protein ratio) analysis showed that only 36 of 1235 statistically relevant proteins, among which 189 are NatB-substrates, were drastically deregulated in *Naa20*^-/-^ MEFs (**Supplementary Dataset 1**, **Table 2**, and **Fig. 2G**). Almost all upregulated proteins were NatA substrates and, where Nt-acetylation could be measured, no variation was observed (**Table 2**). By contrast, the amino terminal sequence of 10 of 21 strongly downregulated proteins indicated that they were predicted as NatB substrates, including ANP32B (Q9EST5), a multifunction protein directly cleaved by caspase-3 and a negative regulator of caspase-3-dependent apoptosis (**Supplementary Dataset 1** and **Table 2**). Although, Nt-acetylation of procaspase-3, -8, and -9 was not quantifiable, caspase-3 and the UBR4 E3-ligase were also found to be reduced in *Naa20*^-/-^ MEFs to some extent (**Supplementary Dataset 1** and **Table 1**). We obtained information of the Nt-acetylation yield of 12 out of the 36 most drastically deregulated proteins. Of 9 downregulated proteins, Nt-acetylation yields were unmodified for five of them (% Nt-acetylation difference <1.3), while four including three NatB substrates - spermidine synthase, caveolae-associated protein 1, and reticulon-4 - displayed a parallel reduction in Nt-acetylation and protein accumulation (**Table 2**).

**Table 2.** Differentially accumulated proteins when the NatB catalytic subunit is inactivated in MEFs. Most affected proteins (2<LFQ protein ratio <0.5) are presented. For an exhaustive list, see Supplementary Dataset1. Blue line divides the downregulated proteins from the upregulated proteins. See also Volcano plot, Fig. 2F.

| Plot# | Entry Name | Protein Description | LFQ Protein Ratio<br>*Naa20<sup>-/-</sup>* / WT | %NTA<br>Difference<br>*Naa20<sup>-/-</sup>* - WT |
| --- | --- | --- | --- | --- |
| 1 | TAGL_MOUSE | Transgelin | 0.331 | -1.3 (Ala2) |
| 2 | SERC_MOUSE | Phosphoserine aminotransferase | 0.492 | -0.4 (iMet) |
| 3 | LYOX_MOUSE | Protein-lysine 6-oxidase | 0.228 | - |
| 4 | USO1_MOUSE | General vesicular transport factor p115 | 0.367 | - |
| 5 | SPEE_MOUSE | Spermidine synthase | 0.444 | -3.3 (iMet) |
| 6 | AN32B_MOUSE | Acidic leucine-rich nuclear phosphoprotein 32 family member B | 0.404 | - |
| 7 | SET_MOUSE | Protein SET | 0.314 | - |
| 8 | CAVN1_MOUSE | Caveolae-associated protein 1 | 0.425 | -15.9 (iMet) |
| 9 | FACR1_MOUSE | Fatty acyl-CoA reductase 1 | 0.474 | - |
| 10 | HNRL2_MOUSE | Heterogeneous nuclear ribonucleoprotein U-like protein 2 | 0.282 | - |
| 11 | LOXL1_MOUSE | Lysyl oxidase homolog 1 | 0.423 | - |
| 12 | RTN4_MOUSE | Reticulon-4 | 0.462 | -4.0 (iMet) |
| 13 | RAB18_MOUSE | Ras-related protein Rab-18 | 0.468 | - |
| 14 | RS30_MOUSE | 40S ribosomal protein S30 | 0.372 | 0.0 (Lys1) |
| 15 | IMA7_MOUSE | Importin subunit alpha-7 | 0.416 | -0.4 (iMet) |
| 16 | EF1B_MOUSE | Elongation factor 1-beta | 0.405 | - |
| 17 | U2AF1_MOUSE | Splicing factor U2AF 35 kDa subunit | 0.450 | - |
| 18 | RLA1_MOUSE | 60S acidic ribosomal protein P1 | 0.406 | -0.1 (Ala2) |
| 19 | YTHD2_MOUSE | YTH domain-containing family protein 2 | 0.438 | -0.2 (Ser1) |
| 20 | QKI_MOUSE | Protein quaking | 0.460 | - |
| 21 | LYAR_MOUSE | Cell growth-regulating nucleolar protein | 0.466 | - |
| 22 | KCD12_MOUSE | BTB/POZ domain-containing protein KCTD12 | 5.143 | - |
| 23 | BGLR_MOUSE | Beta-glucuronidase | 2.274 | - |
| 24 | ASAH1_MOUSE | Acid ceramidase | 2.084 | - |
| 25 | NEP_MOUSE | Neprilysin | 3.020 | - |
| 26 | HYEP_MOUSE | Epoxide hydrolase 1 | 3.681 | - |
| 27 | HXK1_MOUSE | Hexokinase-1 | 3.404 | - |
| 28 | SCOT1_MOUSE | Succinyl-CoA:3-ketoacid coenzyme A transferase 1 | 2.209 | - |
| 29 | TPP1_MOUSE | Tripeptidyl-peptidase 1 | 2.099 | - |
| 30 | INF2_MOUSE | Inverted formin-2 | 2.108 | -0.2 (Ser1) |
| 31 | CSN3_MOUSE | COP9 signalosome complex subunit 3 | 2.273 | 0.0 (Ala2) |
| 32 | S61A1_MOUSE | Protein transport protein Sec61 subunit alpha isoform 1 | 2.887 | -0.3 (Ala2) |
| 33 | H32_MOUSE | Histone H3.2 | 3.601 | - |
| 34 | DDX3Y_MOUSE | ATP-dependent RNA helicase DDX3Y | 2.001 | - |
| 35 | HEXB_MOUSE | Beta-hexosaminidase subunit beta | 2.052 | - |
| 36 | BACH_MOUSE | Cytosolic acyl coenzyme A thioester hydrolase | 2.098 | - |

Taken together, our proteomic investigation suggest that NatB is not likely involved in the proteostasis of all of its substrates, but rather indicates that NatB-mediated Nt-acetylation may still protect a specific set of proteins from degradation, such as components of the apoptotic pathways.

### *Naa20* inactivation reduces the expression of several components of the apoptotic pathways decreasing activation of procaspases in response to TNF-α plus SMAC mimetic and etoposide

Although a significant number of key components of the apoptotic pathway are NatB substrates (see above and **Supplementary** Fig. 1), our proteomic analysis did not allow us to simultaneously quantify the Nt-acetylation yield and protein accumulation of none of them. As it has been shown that Cre recombinase expression could be cytotoxic through unspecific activities^39^, we first confirmed that after 6 days infection it did not affect the expression levels of intrinsic and extrinsic apoptotic components, including the NatB substrates BID and procaspase-8,-9 and -3, independently of *Naa20* inactivation (**Supplementary** Fig. 2). We then performed a targeted time-course assay after *Naa20* inactivation to analyze their accumulation. *Naa20*^-/-^ MEFs showed no variation in apoptotic NatA substrates such as procaspase-6 and SMAC (or DIABLO) (**Fig. 3A**), whose N-termini start with a Thr and Ala, respectively. However, the levels of procaspases-8, -9, -3 and BID significantly decreased in *Naa20*^-/-^ MEFs (**Fig. 3A** and **Supplementary** Fig. 3A), while no differences in *procaspase-8, - 9, -3* and *Bid* transcripts were observed six days after infection (**Fig. 3B**). Protein levels of BAX and APAF1, two other NatB substrates, were unaffected and accumulated in the absence of NAA20 (**Fig. 3A**). Interestingly, both *Bax* and *Apaf1* mRNA levels were markedly upregulated six days post-infection in the absence of NAA20 (**Fig. 3B**), suggesting that BAX protein may be less stable in the *Naa20*^-/-^ background, and that transcriptional upregulation compensates for its decreased stability. Interestingly, we observed the presence of cleaved PARP in the absence of NAA20, despite not detecting active caspase-3 (**Fig. 3A** and **Supplementary** Fig. 3A). This suggests the presence of apoptotic cells in the context of NAA20 deficiency. These data clearly show a strong negative effect on procaspase-3, -8, and -9 and BID expression of NatB inactivation as a result of a reduced Nt-acetylation.

**Fig. 3.**
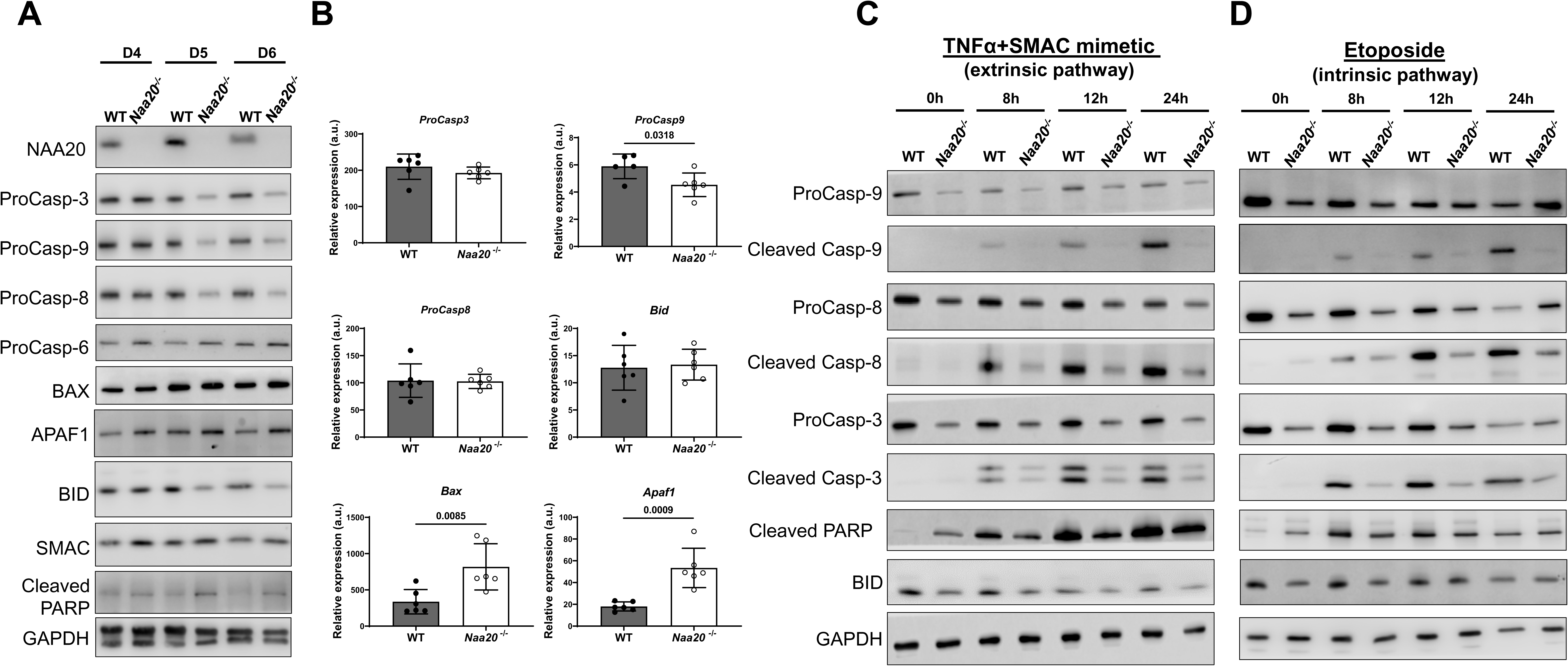
Inactivation of *Naa20* decreases the protein expression of procaspase-8, -9, and - 3, and BID, which block the extrinsic and intrinsic apoptotic pathways. (**A**) Representative western blot images of the NatB substrates procaspase-8, -9, and -3 and of BAX, BID, APAF1 and non-NatB substrate procaspase-6, SMAC and cleaved PARP. MEFs were harvested 4, 5, and 6 days after infection with AdEmpty (WT) or Ad5CMVCre (^-/-^) inducing *Naa20* inactivation. Procaspase-6 and SMAC, which are not NatB substrates, were used as controls. GAPDH was used a loading control. **(B)** Relative quantification by RT-PCR of the same NatB substrates as in (A), six days post-infection. Data are the mean ± SEM of six independent experiments and are normalized to a housekeeping gene (Histone 3). Student’s *t*-test was used to evaluate differences between groups with the obtained values indicated when statistical significance was achieved (p<0.05). **(C)** Representative western blot images of procaspase-8, -9, and -3 and of respective cleaved caspases, cleaved PARP and BID in *Naa20*^-/-^ MEFs six days after infection with AdEmpty (WT) or Ad5CMVCre (^-/-^) 0, 8, 12, and 24 h after treatment with 100 ng/mL TNF-α plus 500 nM SMAC mimetic. **(D)** As in (C) but after treatment with 50 µM etoposide.

The decrease in procaspases accumulation prompted us to assess their activation in *Naa20*^-/-^ MEFs in response to apoptotic inducers like etoposide and TNF-α plus SMAC mimetic. For both stimuli, procaspase-8, -9, and -3 levels slightly decreased in WT cells through proteolytic processing **(Fig. 3C, D,** and **Supplementary** Fig. 3B, C). However, procaspases and BID protein levels in *Naa20*^-/-^ MEFs - while lower at time 0 compared with WT MEFs - remained constant throughout the experiment and, as expected, this decrease was associated with lower levels of cleaved caspases and PARP. BID protein levels in *Naa20*^-/-^ MEFs followed a similar pattern to procaspases.

We next questioned whether the observed blockade of procaspase activation on *Naa20* inactivation could be extended to other apoptosis inducers including tunicamycin or MG132, which induce apoptosis by activating caspase-9 and caspase-8, respectively^40^. Throughout the experiment, the differences in protein levels of cleaved caspase-8, -9, -3, and PARP between *Naa20*^-/-^ MEF and WT cells were diminished for both treatments, in contrast to the significant blockade observed when cells were treated with etoposide and TNF-α plus SMAC mimetic (**Supplementary** Fig. 4A, B, 5A and **B**).

As *Naa20* inactivation in MEFs decreased procaspase activation in response to the intrinsic apoptosis inducer etoposide, we addressed whether blockage of apoptosis could be due to the impairment of BAX activation. There were no differences in mitochondrial BAX and no detectable cytosolic cytochrome *c* (cyt *c*) and SMAC in WT and *Naa20*^-/-^ MEFs under basal conditions (**Fig. 4A** and **Supplementary** Fig. 6A). Curiously, *Naa20*^-/-^ and WT MEFs, 12 h after etoposide treatment, showed similar BAX translocation from the cytosol to mitochondria. However, mitochondrial release of cyt *c* and SMAC was lower in *Naa20*^-/-^ than WT MEFs despite increases in cellular cyt *c* and SMAC after etoposide treatment, as previously described^41^ (**Fig. 4B** and **Supplementary** Fig. 6B). On the other hand, similar cyt *c* and SMAC release from mitochondria to the cytosol, and consequently of procaspase-3 activation were observed in WT and *Naa20*^-/-^ MEFs 8 h after MG132 treatment (**Fig. 4C** and **Supplementary** Fig. 6C).

**Fig. 4.**
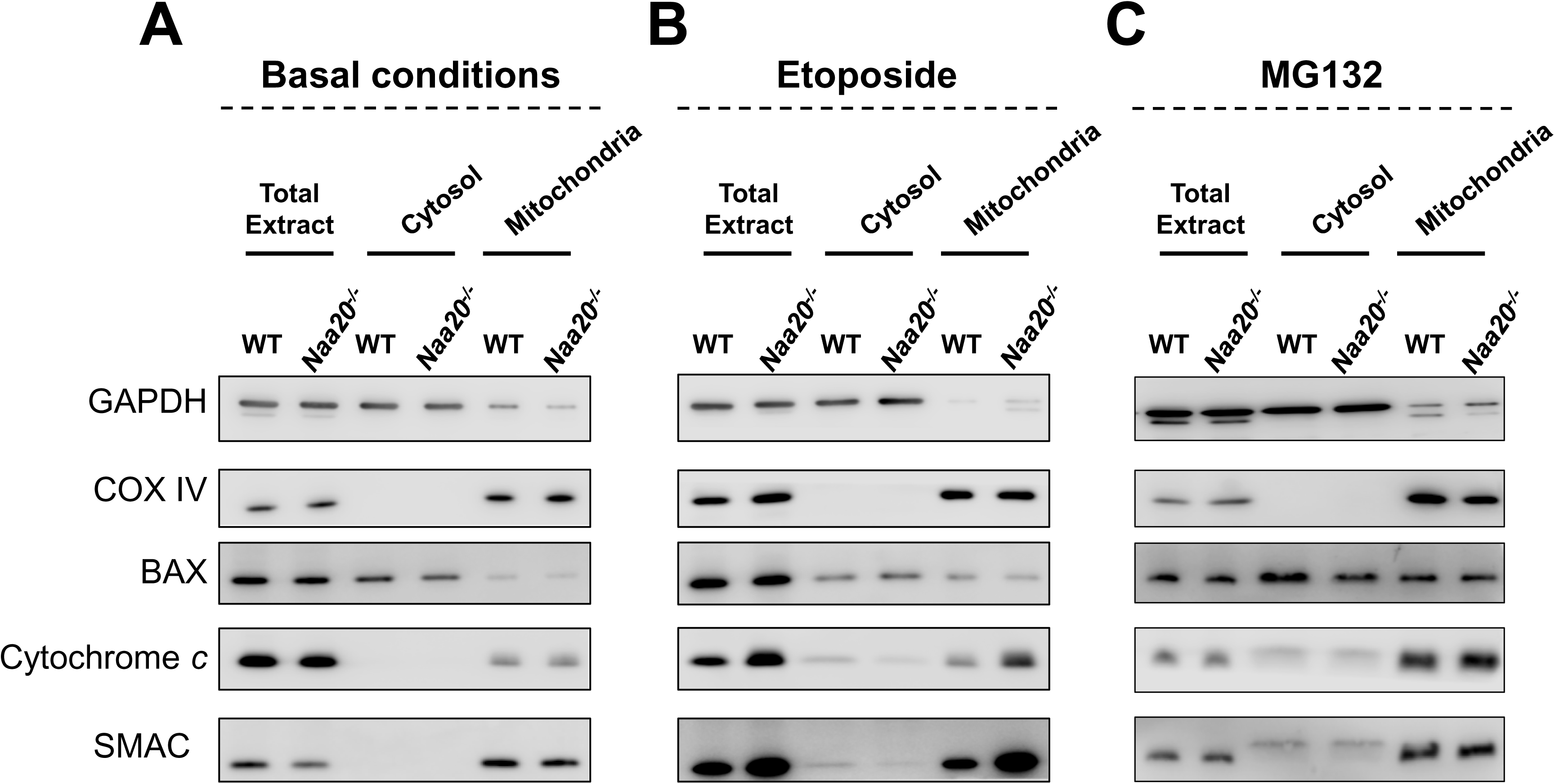
Inactivation of *Naa20* has no impact on BAX localization in response to etoposide, TNF-α + SMAC mimetic, and MG132 treatments but decreases the release of cytochrome *c* and SMAC in response to etoposide. **(A)** Representative western blot images of BAX, cytochrome *c*, COX IV, and SMAC in total extracts, the cytosolic fraction, and mitochondrial fraction of *Naa20*^-/-^ MEFs six days after infection with AdEmpty (WT) or Ad5CMVCre (^-/-^), under basal conditions. **(B)** As in (A) 12 h after 50 µM etoposide treatment. **(C)** As in (A) 8 h after 5 µM MG132 treatment. WT and *Naa20*^-/-^ MEF cells were fractionated 6 days after adenovirus inoculation. Cytosolic GAPDH and mitochondrial COX IV were used as loading controls of cytosolic and mitochondrial fractions, respectively.

### The Arg/N-degron pathway is involved in the reduction of procaspase-8, -9 and BID levels and defective procaspase activation in *Naa20***^-/-^** MEFs

As BID and procaspase-8, -9, and -3 protein levels decreased in the absence of *Naa20* (**Fig. 3A**), we next assessed if the possible lack of the N-terminal acetyl group in NatB substrates was sensed as an Arg/N-degron and consequently targeted for degradation. To this end, we used siRNAs to silence E3 ubiquitin ligases of the Arg/N-degron pathway (*Ubr1*, *Ubr2* and *Ubr4*). As it has been reported a functional complementarity between Arg/N-degron and Ac/N-degron pathways, we also used siRNAs to silence the Ac/N-recognins *Cnot4*, *March6* in *Naa20*^-/-^ and WT MEFs, and evaluated procaspase and BID protein levels (**Fig. 5A** and **Supplementary** Fig. 7A).

**Fig. 5.**
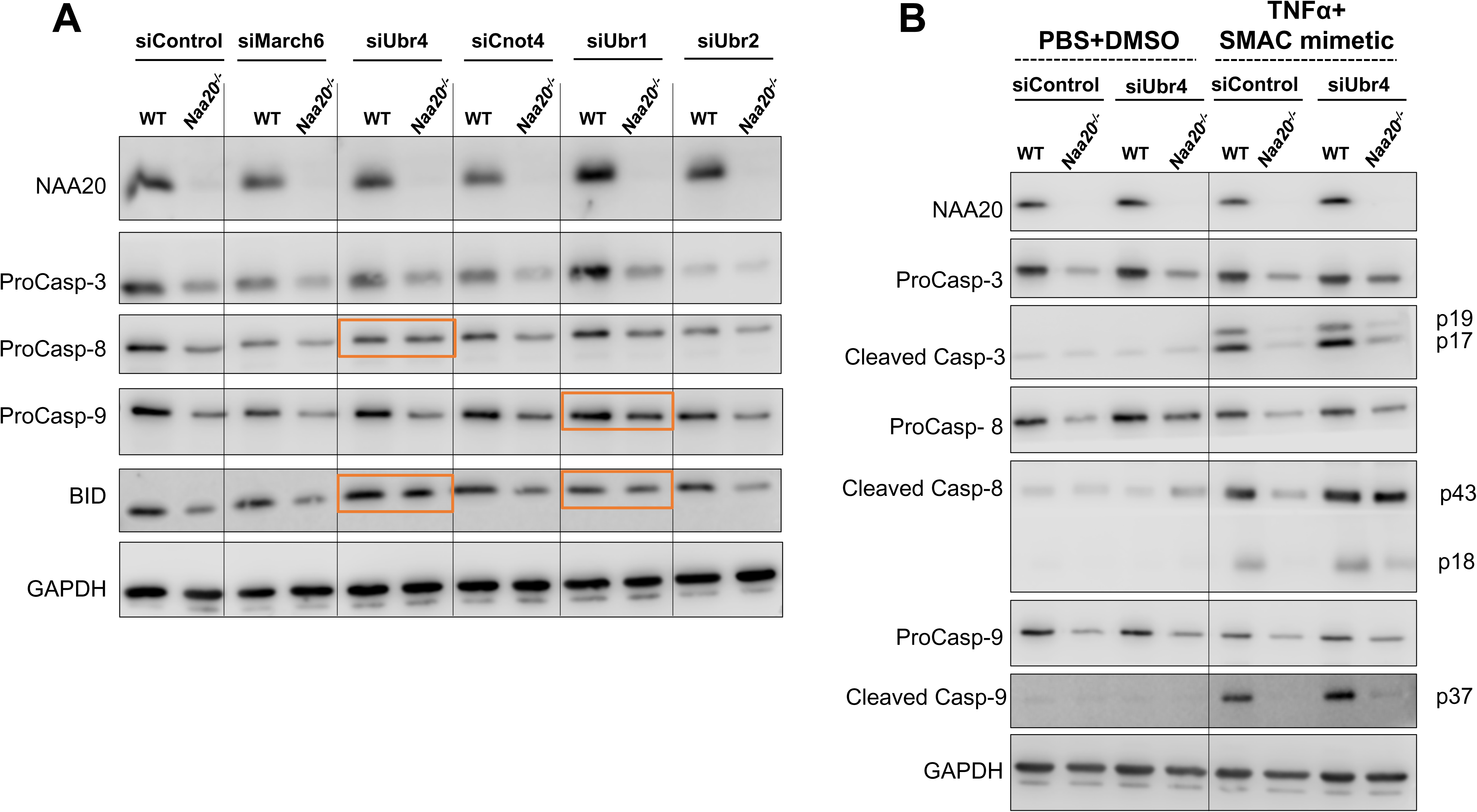
Silencing the ubiquitin ligases *Ubr4* and *Ubr1* in *Naa20*^-/-^ MEFs rescues the decrease in procaspase-8 and -9 protein expression, respectively, as well as BID and promotes the activation of procaspase-8 and procaspase-3 in response to TNF-α plus SMAC mimetic. **(A)** Representative western blot images of procaspase-8, -9, and -3 and BID protein levels after silencing *March6*, *Ubr4*, *Cnot4*, *Ubr1*, and *Ubr2* ubiquitin ligases in *Naa20*^-/-^ MEFs six days after infection with AdEmpty (WT) or Ad5CMVCre (^-/-^). A specific siRNA (siControl) was used as control. **(B)** Representative western blot images of the procaspases and cleaved caspase-8, -9, and -3 protein levels after silencing the *Ubr4* ubiquitin ligase in NAA20 MEFs six days after infection with AdEmpty (WT) or Ad5CMVCre (^-/-^), before (basal conditions) and 12 h after treatment with 100 ng/ml TNF-α plus 500 mM SMAC mimetic. PBS plus DMSO was used as a negative control of TNF-α plus SMAC mimetic. A non-specific siRNA (siControl) was used for control.

While procaspase-3 levels were not positively affected by silencing any of the tested E3 ubiquitin ligases, *Ubr4* silencing increased procaspase-8 levels almost comparable to the level in WT MEFs, while *Ubr1* silencing slightly increased procaspase-9 in *Naa20*^-/-^ MEFs. Moreover, silencing *Ubr4* or *Ubr1* also increased BID protein levels. Interestingly, downregulation of *Ubr2* negatively affected procaspase-8 and procaspase-3 expression in WT MEFs to similar levels as in *Naa20*^-/-^ MEFs. Furthermore, silencing of *March6* and *Cnot4* did not affect the expression levels of procaspases and BID.

### *Ubr4* silencing in *Naa20*^-/-^ MEFs partially reverts the decrease of procaspase-8 expression and its activation in response to TNF-α plus SMAC mimetic

Our data suggest that the procaspase-8, -9 and BID N-termini may be recognized as degradation signals when *Naa20* is inactivated. We, then, wanted to explore the physiological relevance of these NAA20-dependent degradation signals on the induction of the two major apoptotic pathways (e.g., extrinsic and intrinsic ones). We, therefore, investigated whether the procaspase-8 and BID increase observed after silencing of *Ubr4,* and the increase in procaspase-9 and BID after silencing of *Ubr1*, could rescue procaspase activation in *Naa20*^-/-^ MEFs treated with TNF-α plus SMAC mimetic or etoposide, respectively. In addition to restoring procaspase-8 accumulation in *Naa20*^-/-^ MEFs, *Ubr4* silencing also activated caspase-8 12 hours after TNF-α plus SMAC treatment, as it recovered and increased caspase-8 p43 polypeptide production and caspase-8 p18 polypeptide accumulation, respectively (**Fig. 5B** and **Supplementary** Fig. 7B). *Ubr4* downregulation also promoted slight caspase-3 activation in *Naa20*^-/-^ MEFs. Nonetheless, procaspase-3 activation was less pronounced than procaspase-8 activation. Therefore, the decrease in procaspase and caspase-8 in response to TNF-α plus SMAC mimetic caused by *Naa20* inactivation can be partially reverted by silencing the *Ubr4* E3 ubiquitin protein ligase. Conversely, although *Ubr1* silencing restored procaspase-9 accumulation in *Naa20*^-/-^ MEFs (**Fig. 5A**), etoposide treatment barely induced procaspase-9 activation in the double mutant (**Supplementary** Fig. 8A). Of note, no caspase-9 activation was observed when silencing *Ubr4* in response to etoposide (**Supplementary** Fig. 8B).

## DISCUSSION

By targeting around 21% of the proteins, NatB is the second major contributor to the Nt-acetylome and its inactivation or downregulation affects several cellular processes such as actin cytoskeleton function^10, 20, 21, 42^, stress responses^13, 43^, proliferation^22, 23, 26^, and apoptosis^26, 27^. Although NatB is known to regulate apoptosis in MEFs and tumor cells, the association between NatB and apoptosis regulation and its interdependence with N-degron pathways have not previously been investigated. Curiously, several main components of intrinsic and extrinsic apoptosis pathways are NatB substrates based on their second amino acid identity (**Supplementary** Fig. 1). Accordingly, BAX and procaspase-3 are Nt-acetylated by NatB in human cells^16^. In addition, we recently showed that human BAX acetylation by NatB is essential to maintain BAX in an inactive conformation in the cytosol of MEFs^27^. Thus, to further reveal how apoptosis is modulated by NatB-mediated Nt-acetylation, we explored the effect of *Naa20* inactivation on protein and gene expression of several apoptotic pathway components. We observed a strong and selective decrease in the NatB substrates procaspase-8, -9, and -3 and BID but not procaspase-6 and SMAC, both of which are NatA substrates. Nevertheless, when *Naa20* was inactivated, *Apaf1* and to lesser extent *Bax* mRNAs were upregulated while *procaspase-8*, *-9*, and *-3* and *Bid* were not affected. In *D. melanogaster, Naa20* knockdown decreased protein expression of NatB substrate Drk, while accumulation of its mRNA was unaffected^44^. In sum, decreases in procaspase-8, -9, and -3 and BID proteins in *Naa20*^-/-^ MEFs is not caused by transcriptional repression of their genes or through destabilized mRNA, suggesting a mechanism of selective translational repression or instability of these proteins.

Our proteomic analysis is consistent with NatB not being involved in the overall control of its substrates stability, with the accumulation of most of these proteins being not affected by *Naa20* inactivation. Instead, NatB-dependent NTA of specific proteins - particularly of the pro-apoptotic factors procaspase-8, -9, -3 and BID - prevents these proteins from degradation. Therefore, we addressed whether their reduced levels affected *Naa20*^-/-^ MEFs susceptibility to different apoptotic stimuli. Loss of apical initiators caspase-8 and -9 is known to block extrinsic and intrinsic apoptosis, respectively^45, 46^. Our results revealed a decrease in procaspase-8, -9, and -3 activation in response to etoposide- and TNF-α plus SMAC mimetic-induced apoptosis in *Naa20*^-/-^ but not in WT MEFs, reflecting a negative impact of *Naa20* inactivation on both pathways. Intriguingly, *Naa20*^-/-^ MEFs display basal levels of cleaved PARP (Asp214), which suggests the activation of caspase-3, even though the cleaved caspase-3 peptide remains undetectable. However, this particular state does not promote the induction of apoptosis upon cell exposure to etoposide and TNF-α plus SMAC mimetic treatment suggesting the dysfunction or inhibition of apoptotic signaling components.

To understand whether this apoptosis blockade was related to impaired BAX activation, we assessed its subcellular localization and cyt *c* and SMAC mitochondrial release. In *Naa20*^-/-^ MEFs, 12 h after etoposide treatment, BAX mitochondrial translocation was similar to WT MEFs; however, cyt *c* and SMAC release was attenuated, indicating a potential inability to activate procaspase-9 in response to etoposide. As etoposide is a typical caspase-9-dependent drug^47, 48^, the reduction in native and cleaved procaspase-9, cytosolic cyt *c* and SMAC mediated by *Naa20* inactivation likely affect etoposide’s ability to elicit apoptosis. Conversely, TNF-α, a pleiotropic ligand of TNFR1 and 2, promotes cell survival by activating NF-κB or cell death by activating procaspase-8^49^. The observed decrease in procaspase-8 protein levels in the absence of NatB-mediated Nt-acetylation limits caspase-8 activation which ultimately hampers the apoptotic cascade^50^.

An early study investigating the biological function of the catalytic subunit NAA20 in human revealed increased susceptibility of HeLa cells to MG132 after *Naa20* knockdown^26^. Consistently, our previous study with *Naa25*^-/-^ MEFs also revealed increased susceptibility to MG132^27^. Moreover, another study performed in *caspase-9*^-/-^ MEFs revealed efficient MG132-induced but weak etoposide-induced apoptosis, which was restored by synergizing with active cytosolic SMAC^47^. Interestingly, the reduced initial protein levels of procaspase-8, -9 and -3 and their active forms in *Naa20*^-/-^ compared with WT MEFs did not promote significant differential procaspase activation and PARP1 cleavage in response to MG132 or tunicamycin, neither affected BAX activation nor cyt *c* and SMAC release when proteasome was inhibited. It is possible that inhibition of the proteasome by MG132 could counteract the degradation of the proapoptotic factors in the absence of NatB-mediated acetylation. Furthermore, proteasome inhibition and ER stress induction are two biological processes that can promote apoptosis through diverse molecular pathways. Importantly, these mechanisms can operate independently of apoptosome formation and can utilize either the extrinsic or intrinsic pathway. Together, these data indicate that the effect of *Naa20* inactivation on apoptosis induction, likely due to reductions in procaspase-8, -9 and -3 and BID protein levels, is stimulus-dependent.

Since Nt-acetylation prevents protein degradation and plays a role in maintaining partial proteome stability in MEFs, we sought to identify the N-recognins involved in *Naa20*-mediated degradation of apoptosis effectors. Partial reversion of the decrease in procaspase-8, -9 and BID protein expression levels through *Ubr1* or *Ubr4* downregulation confirmed that their unacetylated N-terminus may be sensed as an N-degron and targeted for degradation. Interestingly, *Ubr4* silencing in *Naa20*^-/-^ MEFs restored procaspase-8 activation after TNF-α plus SMAC mimetic treatment. In response to these stimuli, caspase-3 and -9 activation was less pronounced restored when the *Ubr4* N-recognin was downregulated in *Naa20*^-/-^ MEFs, reflecting that cleavage of BID and procaspase-3 by caspase-8 was insufficient to activate properly the initiator and effector caspases of the intrinsic pathway. Similarly, we show that UBR1 recognizes procaspase-9 as an N-degron, as its decreased protein level in *Naa20*^-/-^ MEFs was also partially restored after *Ubr1* silencing. However, *Ubr1* downregulation in *Naa20*^-/-^ MEFs did not promote procaspase-9 activation in response to etoposide, suggesting that it was insufficient to restore activation of the intrinsic pathway.

Taken together, our data reveal that NatB-dependent acetylation of several apoptotic factors protects them from specific UBR E3 ligases-mediated degradation. This further highlights the impact of NatB and Arg/N-degron pathway interdependence on apoptosis. Most interestingly, absence of NatB-dependent acetylation of procaspase-8 exposed the unacetylated N-terminus to UBR4-mediated degradation, thereby limiting caspase-8 activation upon extrinsic stimuli (**Fig. 6**). Our study supports the relevance of NatB-dependent NTA for cellular proteostasis of key pro-apoptotic factors and consequent apoptosis, paving the way for new therapeutic strategies.

**Fig. 6.**
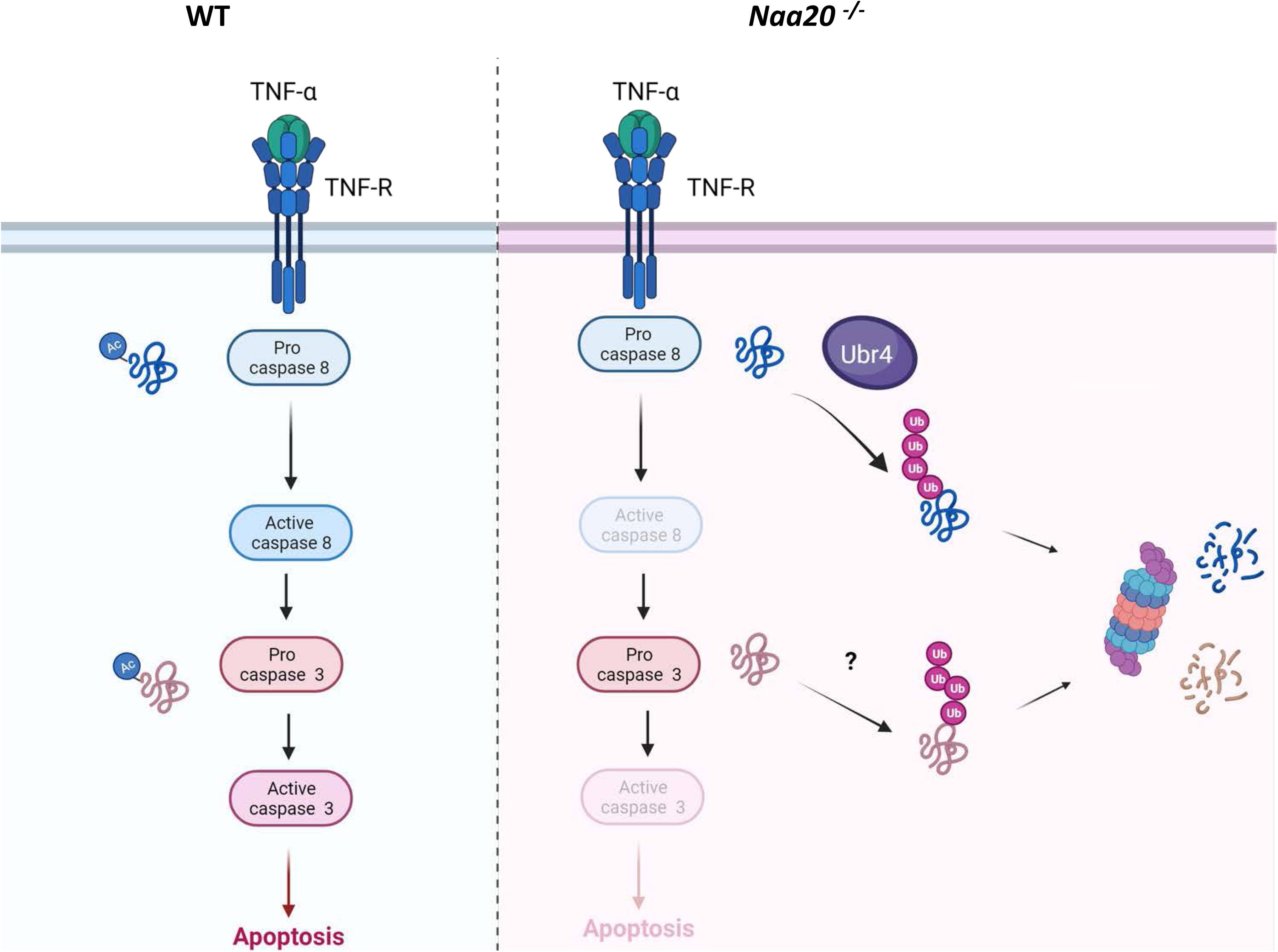
Model depicting the role of NatB-dependent Nt-acetylation in the control of extrinsic apoptosis induction. Procaspase-8 and procaspase-3 are NatB substrates Nt-acetylated in normal cells. TNF-α binding to its receptor in normal cells promotes procaspase-8 activation, which activates procaspase-3 and promotes apoptosis. Inactivation of the NatB catalytic subunit (*Naa20* KO) reduces procaspase-8 Nt-acetylation, and therefore E3-ubiquitin ligase UBR4 recognizes procaspase-8 N-terminus as an N-degron, marks, and sends procaspase-8 for proteasomal degradation. Procaspase-3 is also less stable when *Naa20* is inactivated, but the E3-ubiquitin ligase has not been identified. Therefore, binding of TNF-α to its receptor when NatB is inactivated limits apoptosis induction as a consequence of reduced procaspase-8 and -3 in these cells. Image created with Biorender.

### Data availability

Mass-spectrometry based proteomics data are deposited in ProteomeXchange Consortium (https://proteomecentral.protemeexhange.org) via the PRIDE repository (https://www.ebi.ac.uk/pride/) with the data set identifier PXD029641, username, **Password:** O8uvsKWw.

## Acknowledgements

This work was supported by the KatNat (ERA-NET, ANR-17-CAPS-0001-01) and CanMore (France-Germany PRCI, ANR-20 CE92-0040) grants funded by the French National Research Agency (ANR) to C.G. to support JB.B, by Foundation ARC (ARCPJA32020060002137) grants to T.M., from the facilities and expertise of the I2BC proteomic platform (Proteomic-Gif, SICaPS) supported by IBiSA, Ile de France Region, Plan Cancer, CNRS and Paris-Saclay University, and from ProteoCure COST (European Cooperation in Science and Technology) action CA20113. The proteomic experiments were partially supported by Agence Nationale de la Recherche under projects ProFI (Proteomics French Infrastructure, ANR-10-INBS-08) and GRAL, a program from the Chemistry Biology Health (CBH) Graduate School of University Grenoble Alpes (ANR-17-EURE-0003). Joana P. Guedes acknowledges the PhD fellowship SFRH/BD/132070/2017 funded by FCT.

## Author Contributions

Conceptualization, R.A, C.G. and M.C.R.; Methodology design, C.G., R.A., M.C.R., T.M., JB.B., J.P.G., J.E., and B.C.; Formal analysis, C.G., R.A., M.C.R., T.M., JB.B., J.P.G., J.E., B.C., V.R. and L.S.; Investigation, C.G., R.A., M.C.R., T.M., JB.B., J.P.G., J.E., B.C., V.R. and L.S.; Writing original draft, R.A, C.G. M.C.R. and J.P.G.; Writing-Review and Editing, C.G., R.A., M.C.R., J.P.G., T.M., and JB.B. ; Visualization, C.G., R.A., M.C.R., J.P.G., T.M., and JB.B.; Supervision, R.A, C.G. and M.C.R; Project administration, R.A, C.G. and M.C.R; Funding acquisition, R.A, C.G., and T.M.

## Competing interests

The authors declare no competing interests.

## Materials & Correspondence

Correspondence and material requests should be addressed to Manuela Côrte-Real, Carmela Giglione, and Rafael Aldabe.

## Notes

https://proteomecentral.protemeexhange.org

https://www.ebi.ac.uk/pride/

### Competing Interest Statement

The authors have declared no competing interest.

