## Supplementary material for "NatB-dependent acetylation protects procaspase-8 from UBR4-mediated degradation and is required for full induction of the extrinsic apoptosis pathway"

**Running title:** NatB oversees extrinsic apoptosis

### Supplementary Methods

#### Establishment of mouse embryonic fibroblast strains

MEFs were obtained from *Naa20*$^{tm1a(KOMP)Wtsi}$ transgenic mice generated at the transgenesis unit at the CNB-CSIC using embryonic stem (ES) cells provided by the Knockout Mouse Programme (KOMP). Embryos isolated from the uteruses of pregnant mice were transferred to petri dishes containing sterile PBS. The head and internal organs were removed from embryos. Each embryo was transferred to a well of a six-well plate with trypsin/EDTA, and plates were incubated for 30 min at 37°C. Then, embryos were pulled through an 18-gauge needle to disaggregate them to isolate fibroblasts. Cells were grown in a 100 mm petri dish with Dulbecco's Modified Eagle Medium (DMEM) supplemented with 10% fetal bovine serum (FBS) and 1% penicillin/streptomycin (P/S). When the cells reached confluency, they were re-plated in 60 mm petri dishes and immortalized the next day by retroviral infection with a recombinant virus expressing the SV40 large T antigen and zeocin resistance gene. The infection process was repeated after 24 h to increase efficiency. At confluency, cells were harvested and plated in 100 mm petri dishes. 48 hours later, zeocin (200 µg/mL) was added to the plates for selection of T antigen-expressing cells. Cells were incubated in medium with antibiotic replaced every two days to remove dead cells and refresh the antibiotic. When cell clones were clearly observed, they were harvested with a micropipette and plated in 96-well plates to be grown separately. Each clone was amplified and, once established, they were used in downstream experiments.

#### Real-time PCR

Total RNAs were extracted with the Maxwell® RSC simplyRNA kit (Promega, Madison, WI). Reverse transcription was performed as previously reported[26]. Real-time PCR was performed with the iQ SYBR Green Supermix (Bio-Rad, Hercules, CA) in a CFX96 Real-Time System (Bio-Rad) using specific primers for each gene: Naa20 WT F 5'

CCTTCACCTGCGACGACCTGTT R 5’ GGAATTCAGGGGCGACAGAG; Histone F 5’ AAAGCCGCTCGCAAGAGTGCG R 5’ ACTTGCCTCCTGCAAAGCAC; Caspase-3 F 5’TACATGGGAGCAAGTCAGTGG R 5’CACATCCGTACCAGAGCGAG; Caspase-8 F 5’TCAGAAGAAGTGAGCGAGTTGG R 5’ATCCTCGATCTTCCCCAGCA; Caspase-9 F 5’GGGAAGATCAGGGGACATGC R 5’TCTTGGCAGTCAGGTCGTTC; Apaf1 F 5’CTCCTTGGACGACAGCCATT R 5’AAACACGCGTGGTAAACAGC; Bax F 5’ACCAAGAAGCTGAGCGAGTG R 5’ATGGTTCTGATCAGCTCGGG; and Bid F 5’ GGCGTCTGCGTGGTGATTC R 5’ CCAGTAAGCTTGCACAGGCA. Transcript was quantified using the formula: $2^{ct(\beta-Histone)-ct(gene)}$, with ct being the point at which the fluorescence rises substantially above background fluorescence.

**Immunofluorescence and confocal microscopy**

For immunofluorescence experiments, cells were seeded in six-well plates containing glass coverslips. Six days post-infection, cells were fixed with 4% paraformaldehyde (PFA, 16% formaldehyde solution; Thermo Fisher Scientific, Waltham, MA) for 15 min at room temperature (RT). After rinsing with 1× PBS, cells were permeabilized with 0.1 % Triton X-100 (Sigma-Aldrich, St. Louis, MO) over 15 min at RT and washed. Following this process, cells were incubated with anti-α-vinculin (V9131, Sigma-Aldrich) antibodies and Alexa Fluor 488-phalloidin (Invitrogen, Waltham, MA) in TBS-Tween 20 (TBST 1×)-3 % BSA for 30 min, 37ºC. Subsequently, cells were washed and incubated with anti-mouse IgG Cy3 conjugate developed in sheep (Sigma-Aldrich) or anti-rabbit IgG Alexa488 conjugate developed in donkey (Molecular Probes, Eugene, OR) over 30 min at 37ºC. After mounting the coverslips in Vectashield mounting medium with DAPI (Vector Laboratories, Newark, CA), samples were maintained at 4°C until visualization. Images were acquired with an Axiovert 200M confocal LSM 510 META Zeiss microscope using a 40× objective.

**Protein extraction**

Cells were harvested, and 2× Laemmli Sample Buffer (Bio-Rad) plus β-mercaptoethanol with sodium dodecyl sulfate (SDS) 2% (Bio-Rad) supplemented with Tris-HCl 0.5 M pH 7.4 and protease inhibitors (1 mM PMSF, 0.001 mg/mL aprotinin, 1 mM sodium orthovanadate, and 1 mM sodium pyrophosphate (Roche, Basel, Switzerland)) was used to collect cell lysates. The soluble protein concentration was normalized using the Revert 700 Total Protein Stain (LI-COR) and Image Studio Lite Software (LI-COR, Lincoln, NE).

**Western blotting**

Protein samples were separated by SDS polyacrylamide gel electrophoresis and transferred onto nitrocellulose membranes (Bio-Rad). Next, to avoid non-specific interactions, membranes were blocked in 5% BSA or 5% non-fat milk in 1× TBST for 30 min at RT with agitation. Membranes were then incubated 1 h at RT with primary antibodies, namely mouse monoclonal anti-GAPDH (AbD Serotec, Bio-Rad), mouse monoclonal anti-α-tubulin (Sigma-Aldrich), and rabbit polyclonal anti-Nat5 (ProteinTech). Rabbit polyclonal anti-caspase 3, anti-cleaved caspase 3, anti-cleaved caspase 9, anti-cleaved caspase 8, anti-BAX; rabbit monoclonal anti-caspase 8, anti-Cyt *c*, anti-APAF1, anti-COX IV, anti-caspase-6, anti-SMAC; and mouse monoclonal anti-caspase 9 and anti-cleaved PARP (Asp214) were all purchased from Cell Signaling Technology (Danvers, MA). Rat monoclonal anti-BID (truncated and full length) was purchased from R&D systems. Subsequently, membranes were incubated with secondary anti-mouse IgG or anti-rabbit IgG antibodies (Cell Signaling Technology). Chemiluminescence detection was performed using the ECL Ultra detection system (Lumigen, Southfiled, MI) and an Odyssey® Fc Imaging System (LI-COR).

**Cell proliferation assay**

Cells were seeded in six-well plates and infected the day after. Then, cells were trypsinized and re-plated two and five days after infection, as described above. Cell counting was performed at days 2, 3, 4, 5, and 6 post-infection, and four wells per condition/per day were used. Cells were counted using a TC20™ Automated Cell Counter (Bio-Rad).

**MEF fractionation**

Cells were seeded in six-well plates, infected, and, six days post-infection, cells were harvested (basal conditions) or treated with 50 µM etoposide and 5 µM MG132 for 12 h and 8 h, respectively. Then, 6.6 x $10^6$ cells per condition were harvested and cell fractionation was performed using a Cell Fractionation kit (Abcam, Cambridge, UK; AB109719) following the manufacturer's guidelines. 1 x $10^6$ cells per condition were used for total extracts. 4× Laemmli Sample Buffer (Bio-Rad) plus β-mercaptoethanol with SDS 2% (Bio-Rad) supplemented with Tris-HCl 0.5 M pH 7.4 and protease inhibitors was used to collect cellular fractions. GAPDH and COX IV were used as cytosolic and mitochondrial controls, respectively. Protein bands were quantified Image Studio Lite Software System (LI-COR).

**siRNAs used to silence E3 ubiquitin ligase**

The following siRNAs were used: siRNA Silencer™ Select Negative Control No. 1 (4390843), Silencer® Select Ubr1 siRNA (4390771-s75706), Silencer® Select Ubr2 siRNA (4390771-s104948), Silencer® Select Ubr4 siRNA (4390771-s87462), Silencer® Select C4 siRNA (4390771-s79257), and Silencer® Select March6 siRNA (4390771-s104524), all from Thermo Fisher Scientific.

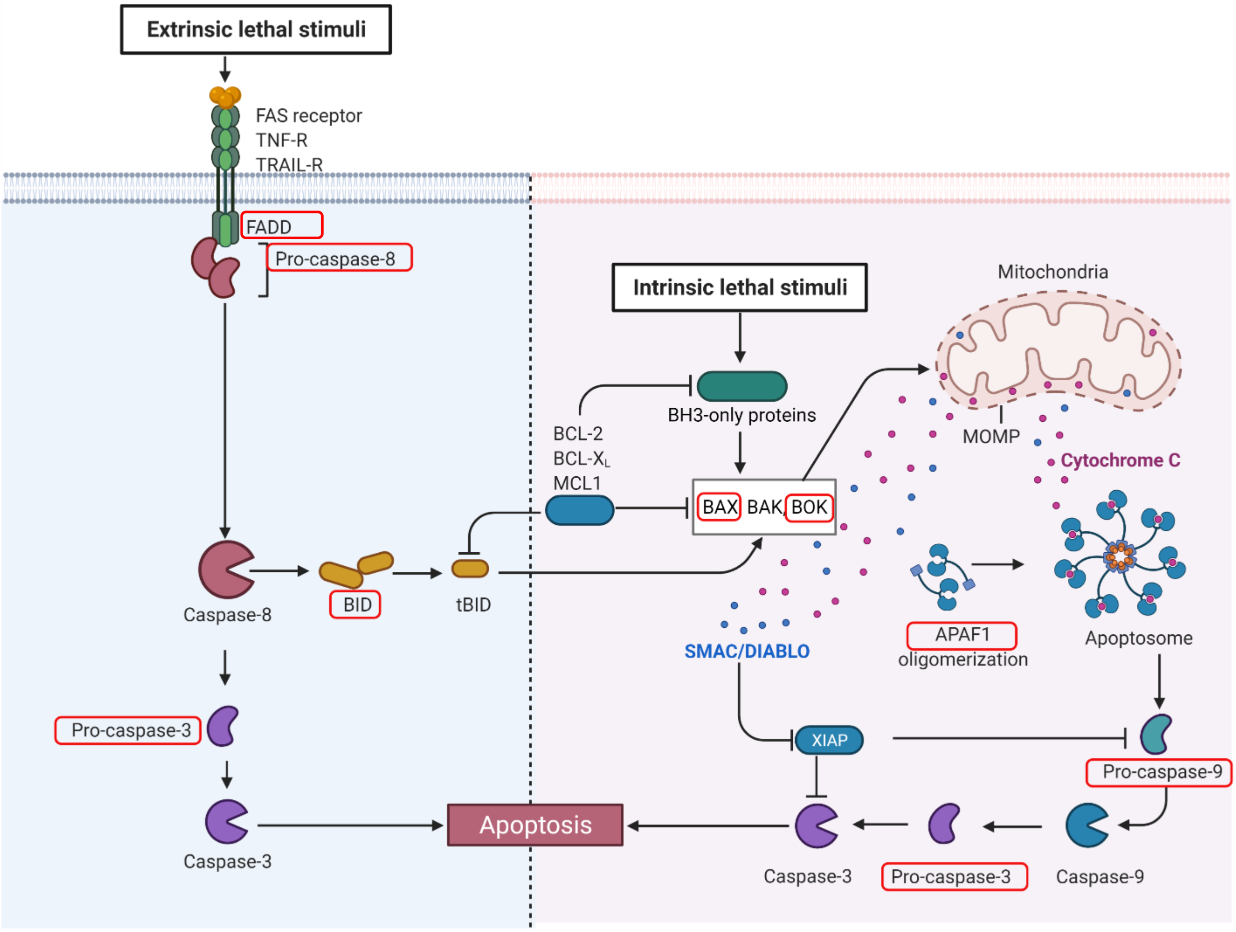


**Figure S1. Intrinsic and extrinsic apoptotic pathways.** Main components of intrinsic and extrinsic apoptotic pathways are indicated. Putative NatB substrates, as they display Asp-,Glut-, Asn- or Gln- at position two, are included in red squares. Binding of extrinsic lethal stimuli like TNF-α and Fas ligand to their receptors promotes caspase-8 activation that in type I cells can initiate apoptosis after caspase-3 proteolytic activation. In type II cells, apoptosis induction requires cleavage of BID into tBID by caspase-8 to initiate the mitochondrial apoptosis pathway. The intrinsic apoptosis pathway is initiated by several intrinsic lethal stimuli (DNA damage, ER stress, replication stress, etc.) that lead to irreversible mitochondrial outer membrane permeabilization (MOMP) controlled by pro-apoptotic (BAX, BAK, BOK) and anti-apoptotic (BCL-2, BCL-XL, MCL1) proteins with BCL2 homology (BH) domains. MOMP directly promotes the cytosolic release of cytochrome *c* (cyt *c*) and SMAC/DIABLO from the mitochondria. When in the cytosol, cytosolic cyt *c* binds to APAF1, promoting its oligomerization and assembly in a supramolecular complex known as apoptosome, which is responsible for activating procaspase-9. Activated caspase-9 can catalyzes the proteolytic activation of procaspase-3 and precipitates apoptosis (Biorender).

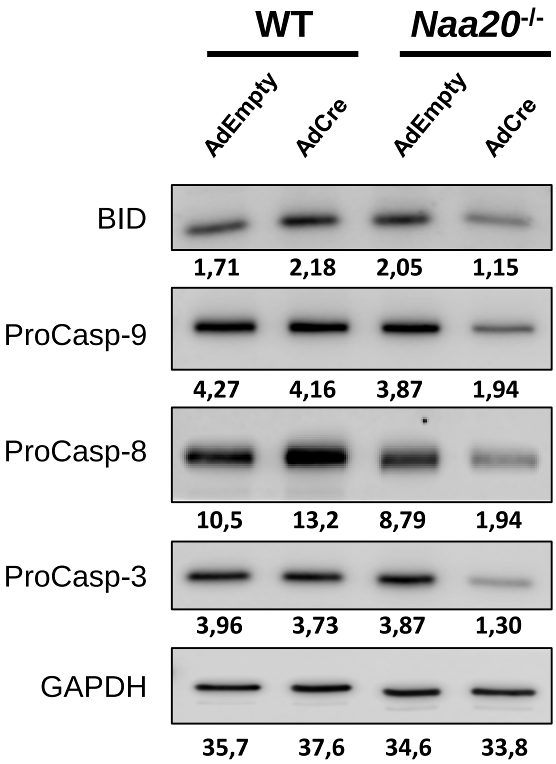


**Figure S2. Effect of Cre recombinase expression on apoptosis components in wild-type and *Naa20*$^{-/-}$ MEFs.** Representative western blot images of BID and the procaspases -8, -9 and -3 protein levels in wild type and *Naa20*$^{-/-}$ MEF cells 6 days after infection with AdEmpty or Ad5CMVCre. Each band intensity value (a.u) is indicated under each band.

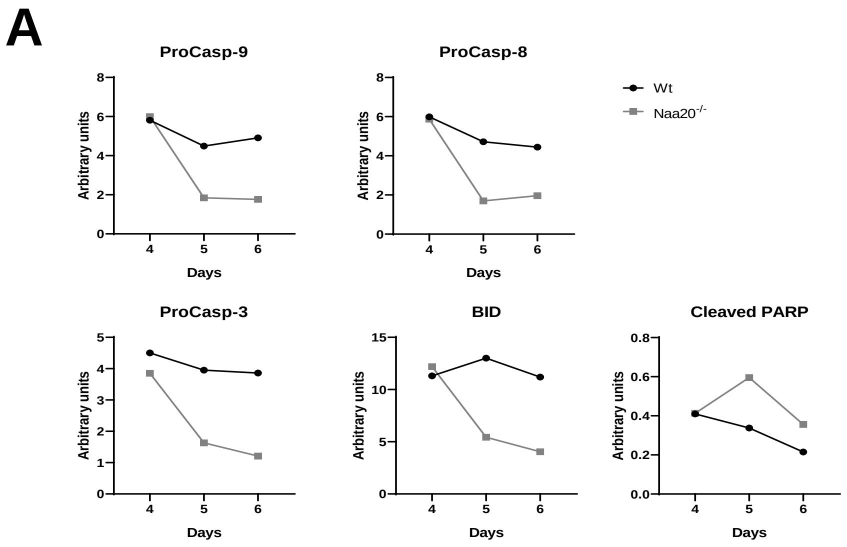


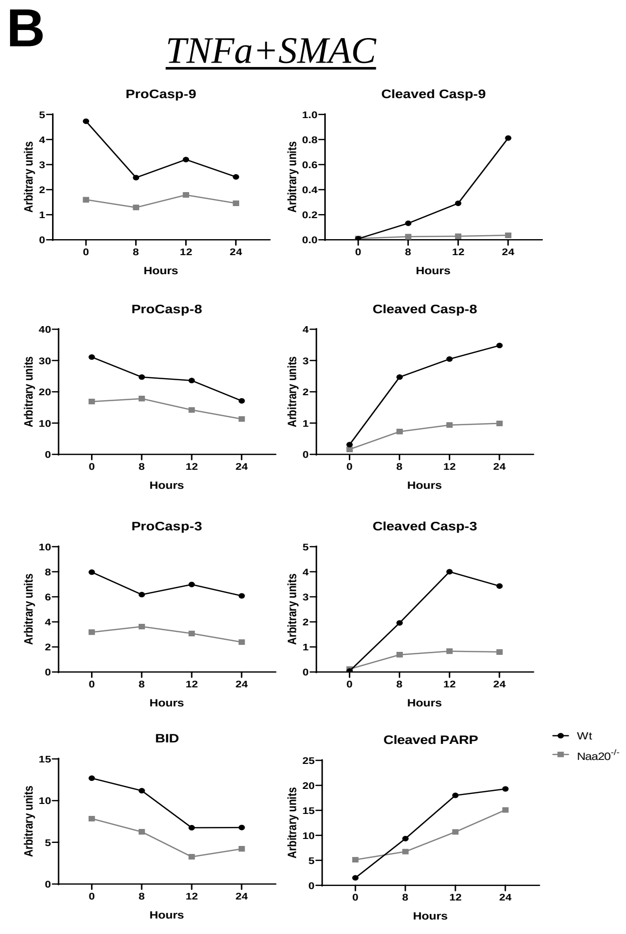


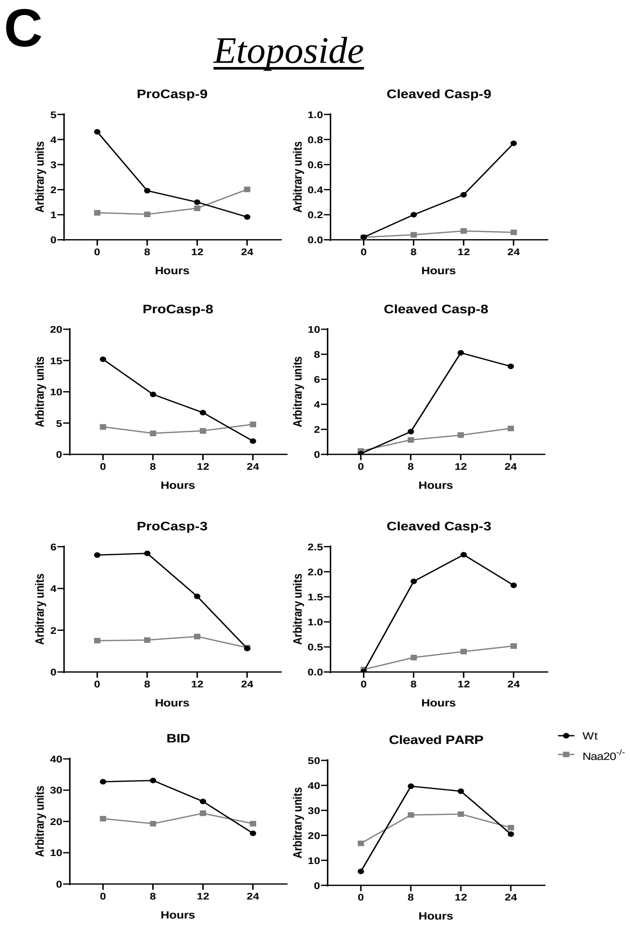


**Figure S3. Quantification of representative western blot signals from Fig. 3.** Panels (A), (B) and (C) correspond to western blot signals from Figs. 3A, 3C and 3D, respectively. Image Studio software was used for the signal quantification.

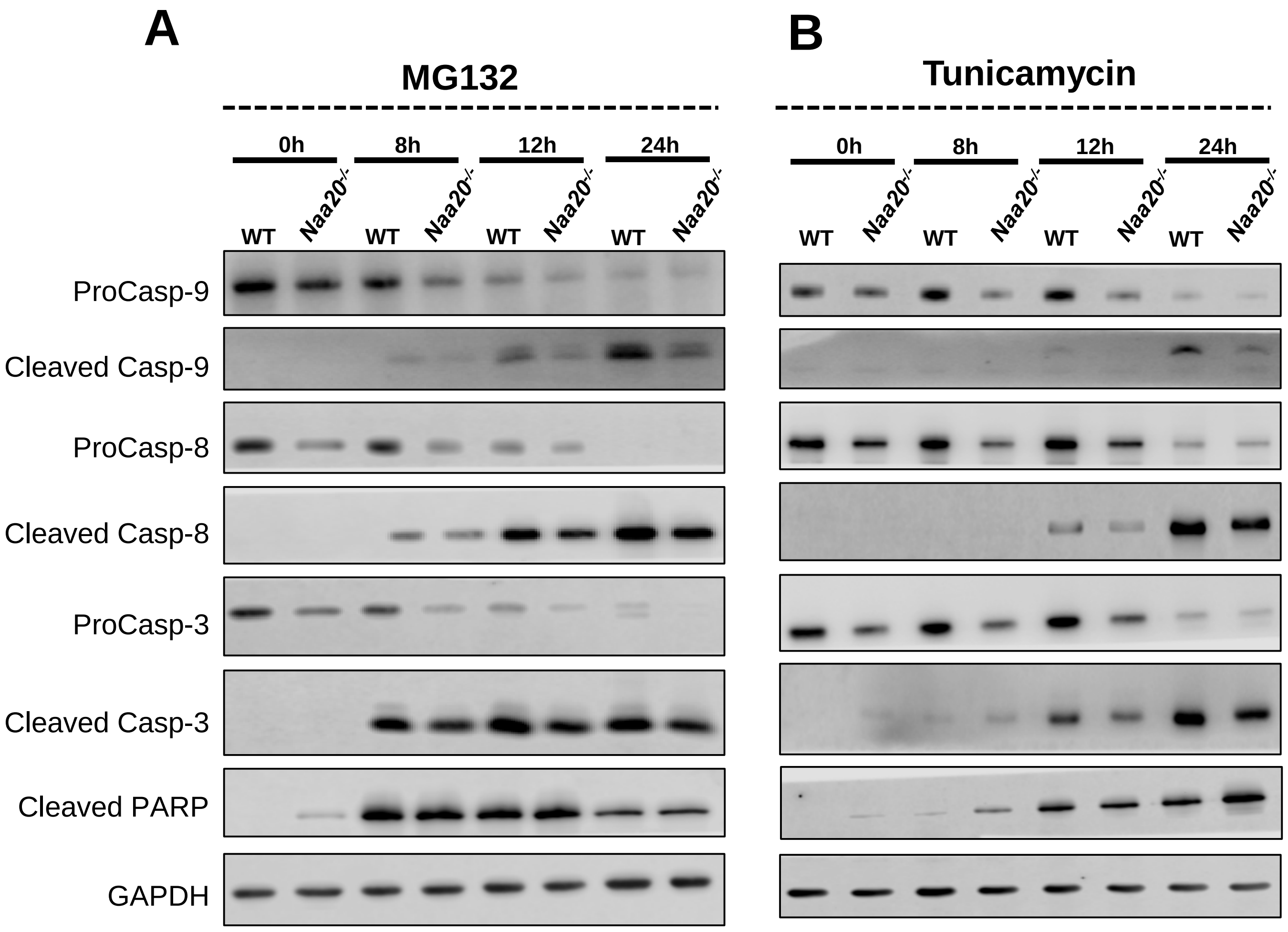


**Figure S4. Inactivation of NAA20 subunit causes a reduction in the expression levels of procaspases but this reduction was attenuated along treatment with MG132 and tunicamycin.** Representative western blot images of the main apoptotic effectors at 0, 8, 12 and 24 h after treatment with (A) 5 µM of MG132 and (B) 5 ng/µl of tunicamycin of control and *Naa20*$^{-/-}$ MEF cells, 6 days after infection with AdEmpty (WT) or Ad5CMVCre (*Naa20*$^{-/-}$).

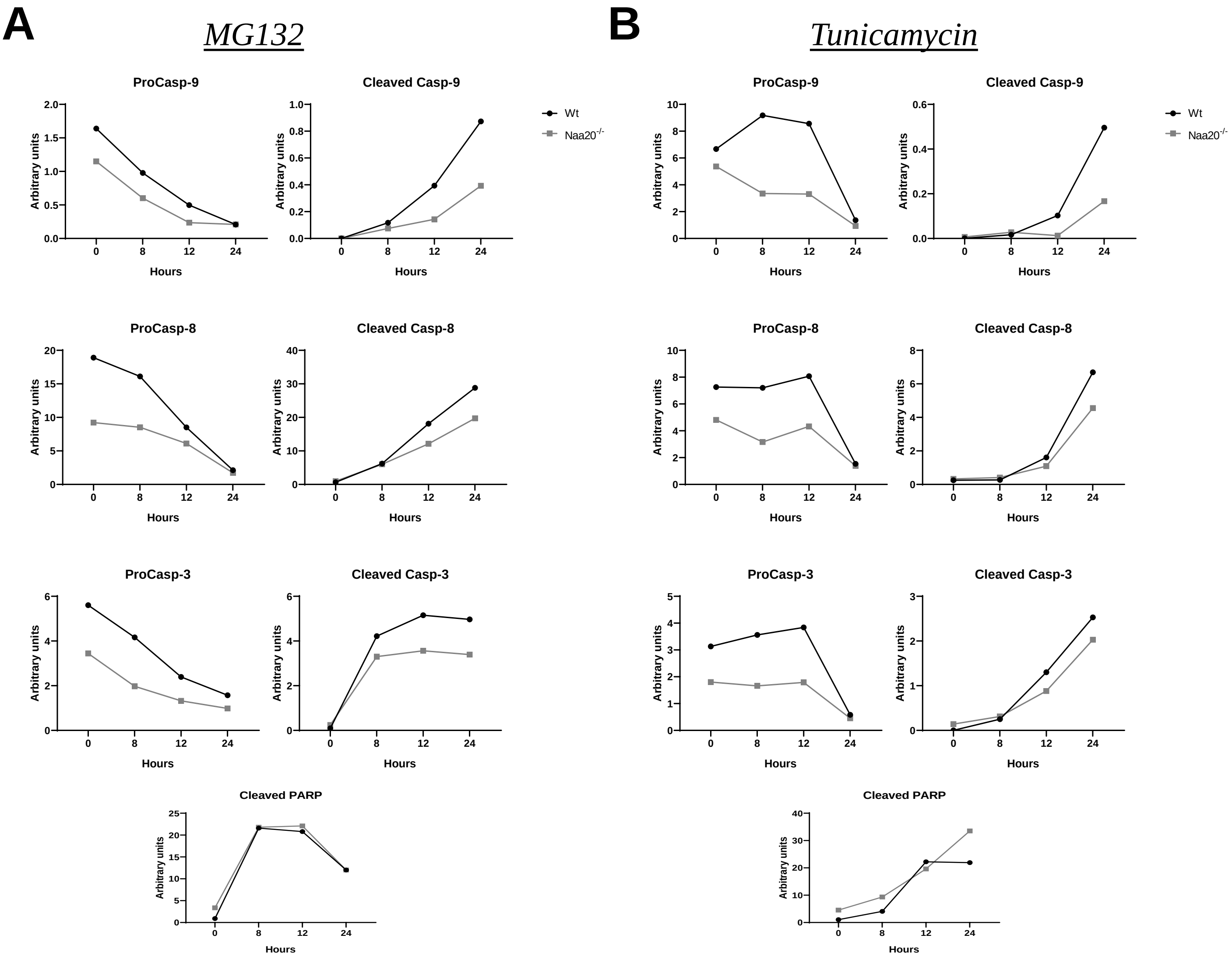


**Figure S5. Quantification of representative western blot signals from Fig. S3.** Panels (A) and (B) correspond to Supplementary Fig. 4A and 4B, respectively. Image Studio software was used for the signal quantification.

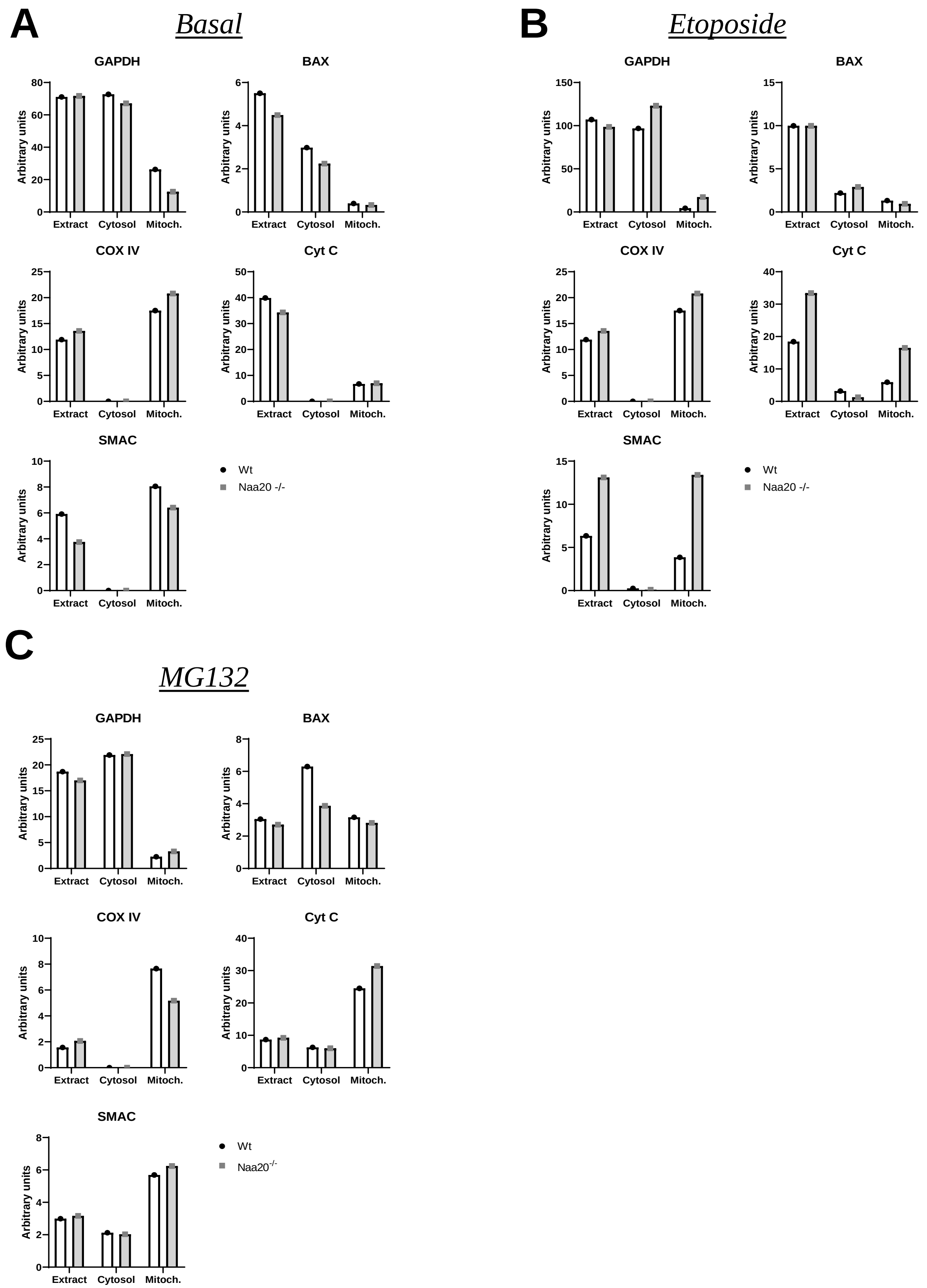


**Figure S6. Quantification of representative western blot signals from Fig. 4.** Panels (A), (B), and (C) correspond to Figs. 4A, 4B, and 4C, respectively. Image Studio software was used for the signal quantification.

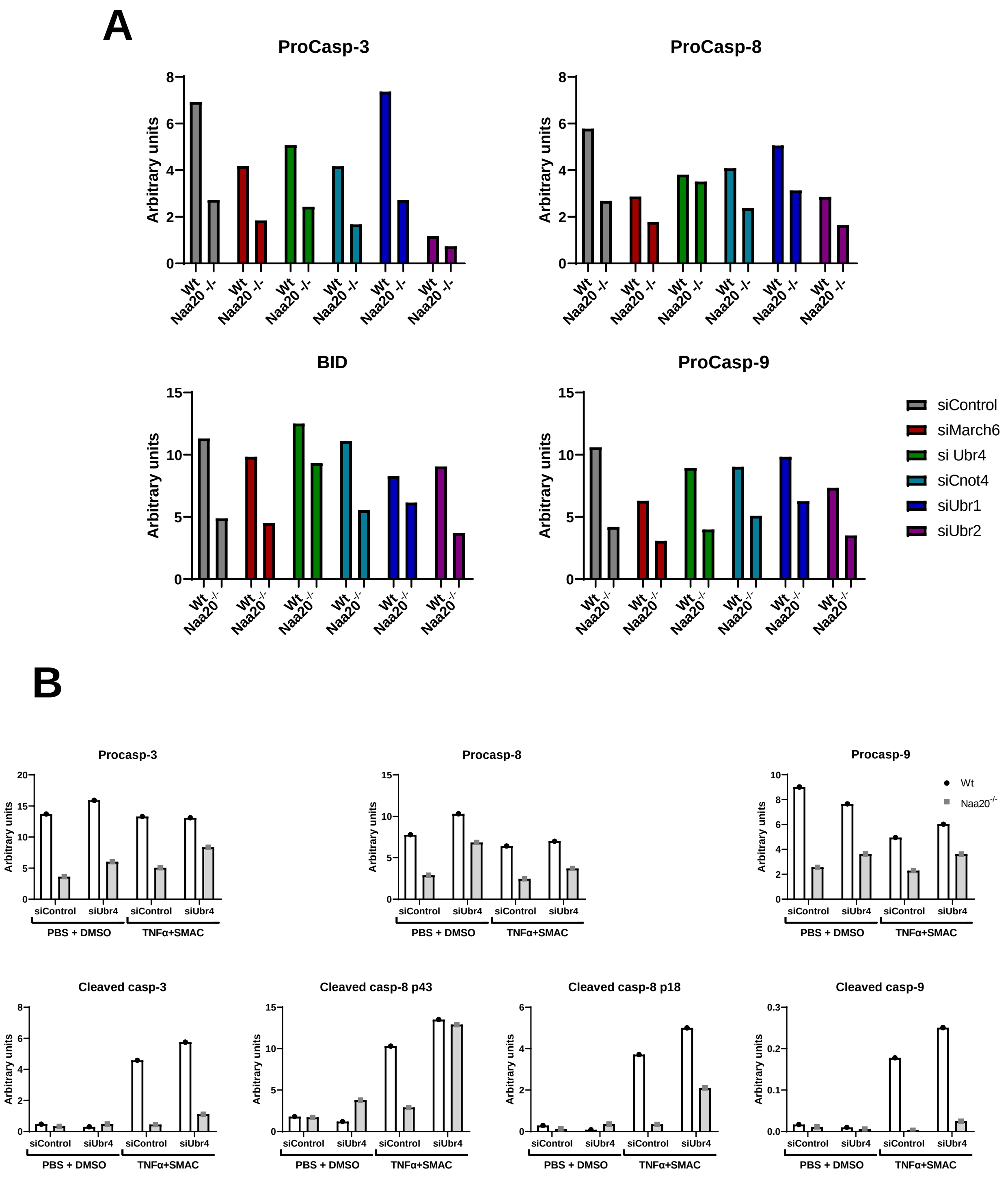


**Figure S7. Quantification of representative western blot signals from Fig. 5. P**anels (A) and (B ) correspond to Fig 5A and B, respectively. Image Studio software was used for the signal quantification.

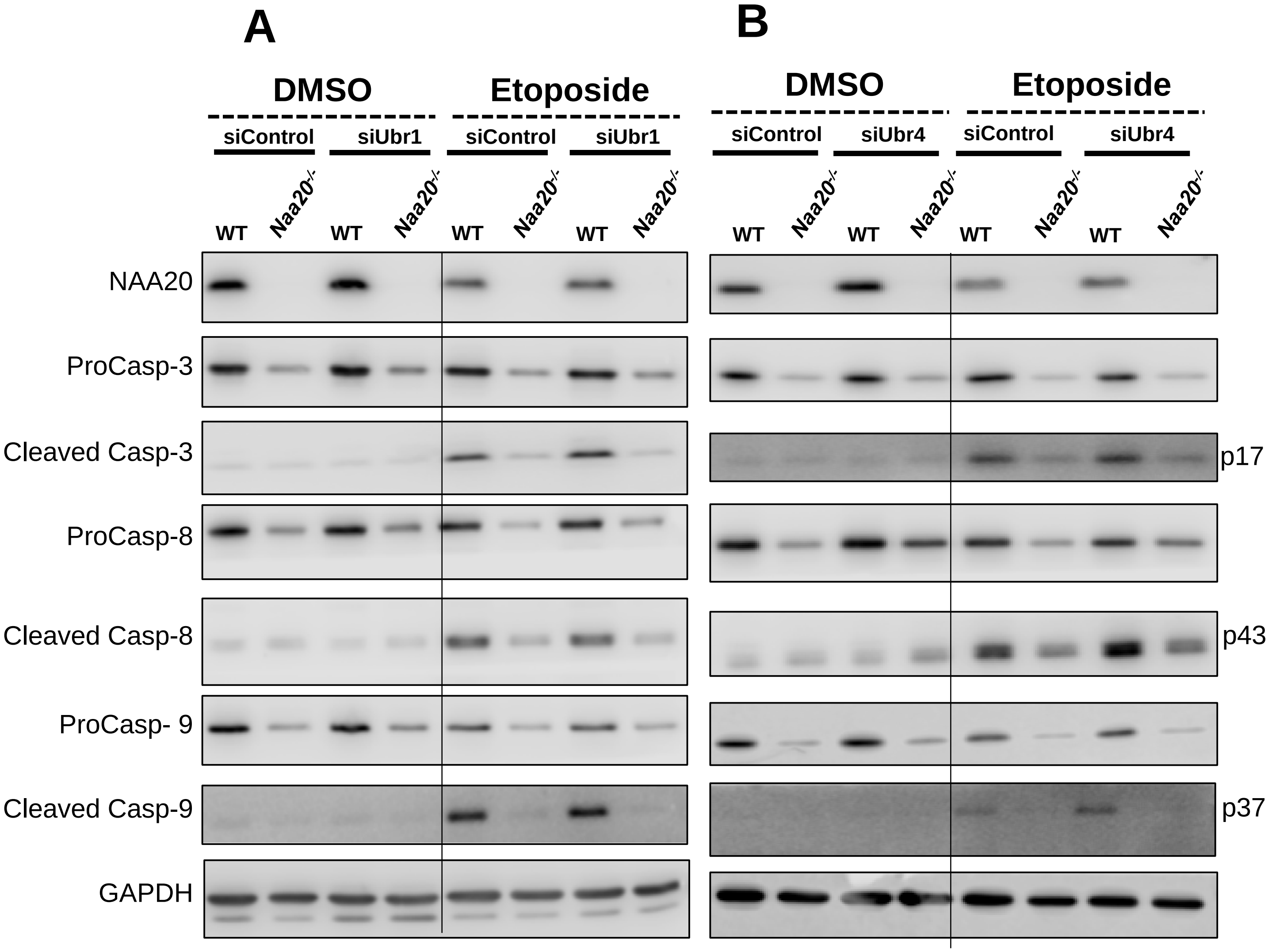


**Figure S8. Silencing of *Ubr4* and *Ubr1* is unable to restore procaspase-9 in response to etoposide in *Naa20*$^{-/-}$ MEF cells.** Representative western blot images of the procaspases and cleaved caspases-8, -9 and -3 protein levels after silencing the *Ubr1* E3 ubiquitin ligase (**A**) or *Ubr4* E3 ubiquitin ligase (**B**) in Naa20 MEF cells 6 days after infection with AdEmpty (WT) or Ad5CMVCre (-/-), before (basal conditions) and 12 h after 50 μM etoposide treatment. Vehicle (DMSO) was used as a control.

| | WT (8 replicates) | *Naa20*$^{-/-}$ (8 replicates) | Unique N-termini |
|---|---|---|---|
| **Identified N-termini** | **13149** | **11899** | |
| **Non-redundant proteoforms** | **1844** | **1675** | **1988** |
| **Quantified N-termini** | **1073** | **945** | **1191** |
| *Full acetylation (NTA > 95%)* | *734* | *586* | - |
| *Partial acetylation* | *90* | *126* | - |
| *No acetylation (NTA < 5%)* | *249* | *233* | - |
| **iMet (pos. 1)** | **244** | **181** | **258** |
| *Full acetylation (NTA > 95%)* | *194* | *99* | - |
| *Partial acetylation* | *25* | *61* | - |
| *No acetylation (NTA < 5%)* | *25* | *21* | - |
| **+NME (pos. 2)** | **688** | **632** | **769** |
| *Full acetylation (NTA > 95%)* | *528* | *478* | - |
| *Partial acetylation* | *55* | *56* | - |
| *No acetylation (NTA < 5%)* | *105* | *98* | - |
| **Processed (pos. > 2)** | **141** | **132** | **164** |
| *Full acetylation (NTA > 95%)* | *12* | *9* | - |
| *Partial acetylation* | *10* | *9* | - |
| *No acetylation (NTA < 5%)* | *119* | *114* | - |

**Supplementary Table 1. Summary of N-terminal profiling of WT and *Naa20*$^{-/-}$ MEF cells.** This table displays the number of N-termini identified and quantified in both conditions. For each sample group, 4 biological replicates were processed, every one of them being analyzed in parallel on two different mass spectrometers, an LTQ-Orbitrap Velos (Thermo Fisher Scientific) and a TIMS-ToF (Bruker), resulting in a combination of 8 replicates per sample. The quantified N-termini were sorted in 4 different categories: their total number, the ones beginning with the initiating methionine (« iMet »), those that underwent N-terminal methionine excision (« +NME ») and the matured N-termini (« Processed »). In each of these categories, the quantified N-termini were either quantified as fully acetylated (NTA yield > 95%), non-acetylated (NTA < 5%), or partially acetylated. Comparison of these numbers allowed to determine the effect of the *Naa20*$^{-/-}$ mutant on the N-terminal acetylation.
